# Global Proteomic Analysis of Colorectal Cancers Stratified by Microsatellite Instability Subtype Reveals Protein Differences

**DOI:** 10.64898/2026.01.28.702312

**Authors:** Fernando Tobias, Emily R. Sekera, Xingzhao Xiong, Fei Fang, Heather Hampel, Rachel Pearlman, Xiaowen Liu, Liangliang Sun, Amanda B. Hummon

**Author notes:** These authors contributed equally to this work (Fernando Tobias, Emily R. Sekera). Corresponding Authors: Liangliang Sun; Xiaowen Liu and Amanda Hummon.

## Abstract

Lynch syndrome, historically known as hereditary nonpolyposis colorectal cancer, is caused by germline mutations in the DNA mismatch repair (MMR) genes, *MLH1, MSH2 (EPCAM), MSH6*, and *PMS2*. While the genetic changes associated with Lynch syndrome have previously been characterized, there have not been studies of the associated proteomic alterations, in part because of the limited availability of primary samples and the absence of *in vitro* model systems. In this study, the first large-scale tissue proteomic assessment of Lynch syndrome samples as well as three other subtypes of colorectal cancer was completed with specimens from the Ohio Colorectal Cancer Prevention Initiative. The cohort contained three groups of microsatellite unstable (MSI-high) colorectal cancer (CRC) patients (Lynch syndrome, double somatic MMR mutation, and MLH1 hypermethylation) and a group of microsatellite stable (MSS) CRC patients. A total of 122 tumor and complimentary normal mucosa samples from 61 patients were evaluated using label-free bottom-up proteomic analysis identifying 7,352 proteins. Hierarchical clustering analysis of the global proteome showed that the MSS group was significantly different than the three MSI-high groups. Of the 1,607 proteins found to be dysregulated within CRC patients, 648 proteins were specific to Lynch syndrome and 260 proteins were dysregulated across all four subtypes studied. Further examination of dysregulated proteins in response to age of diagnosis and patient’s sex assigned at birth were also evaluated. This study fills a critical gap as the first proteomic characterization of Lynch syndrome samples to date. Data are available via ProteomeXchange with identifier PXD073693.

**Teaser:** This is the first study of the global proteomic differences between Lynch syndrome and other forms of colorectal cancer.

**Blurb:** Lynch syndrome is the most common form of hereditary colorectal cancer and causes a genetic predisposition to other cancer types. Diagnosing Lynch Syndrome is extremely useful for treatment of a current cancer as well as future cancer prevention for an individual and their family members. The widespread protein changes that occur in Lynch tumors have never been examined. This study is the first global evaluation of the proteomic changes in the tumors and normal mucosa of Lynch Syndrome patients.

## Introduction

Colorectal cancer (CRC) is the third most common cancer diagnosed worldwide in both men and women.^1^ The majority of CRC cases are microsatellite stable (MSS), with 15% exhibiting high microsatellite instability (MSI-H). MSI is a condition characterized by the accumulation of errors in microsatellite repeat sequences due to a deficient DNA mismatch repair (MMR) system. This deficiency can arise from three primary mechanisms: Lynch syndrome (LS), double somatic MMR gene mutations, and epigenetic silencing of *MLH1* (hypermethylation).

- Lynch syndrome (LS) is the most common form of hereditary CRC and causes a genetic predisposition to other cancer types, including endometrial, ovarian, stomach, and urothelial, among others. Diagnosing LS is extremely useful for treatment implications of a current cancer as well as future cancer prevention for an individual and their family members.
- Tumors caused by “double somatic” MMR mutations are a more recently described phenomenon that involves two separate somatic mutations in the same MMR gene on opposite alleles (*in trans*) within the same cell, leading to MSI in the absence of a germline mutation and typically presenting in a sporadic manner. ^2, 3^
- The most common cause of MSI-H CRC is epigenetic silencing (ES) due to hypermethylation of the *MLH1* promoter which inhibits expression of the MLH1 gene without altering the DNA sequence. ^4^ *MLH1* promoter hypermethylation is typically acquired and only rarely inherited. MSI-H tumors typically have a better prognosis and are more likely to respond to immunotherapies, such as immune checkpoint inhibitors.
- MSS colorectal cancers, characterized by stable microsatellites, harbor an intact MMR system. These tumors tend to follow a different tumorigenesis pathway, marked by chromosomal instability and often involve somatic mutations in genes such as *KRAS* and *TP53*. MSS tumors generally have a poorer prognosis and are less responsive to immunotherapy compared to MSI tumors.

A better understanding of these four types of CRC at the molecular level could prove pivotal in improving early diagnosis and designing therapeutic options. ^5^

Much effort has been devoted to understanding CRC, including MSI tumors, at the genome and transcriptome levels. However, these analyses cannot accurately reflect the protein-level information as post-transcriptional regulation can modulate protein expression and post-translational modifications (PTMs) can influence protein functions. The generation of proteome-level information is imperative for bridging the genotype and phenotype of CRC. Mass spectrometry (MS)-based proteomics has become increasingly popular for high-throughput and large-scale measurement of proteins in cancer cells and tissues for pursuing a better understanding of underlying molecular mechanisms of cancer progression and for discovering new protein biomarkers for more accurate and earlier diagnosis. ^6-13^ However, there are limited proteomics datasets generated from LS samples, most likely due to the limited availability of clinically obtained tissue. To discover the possibility of new protein biomarkers, we performed the first large-scale quantitative proteomics study of LS tissue samples and compared the tissue proteome landscape across different subtypes of CRC: MSS, MSI: LS, MSI: Double Somatic (DS), and MSI: *MLH1* hypermethylation (*MLH1*_hm_), as well as healthy controls (**Table 1** and **Figure 1**). By integrating demographic metadata with quantitative proteomics, this study provides critical insights into the molecular heterogeneity of CRC and the influence of patient-specific factors on tumor biology, setting the stage for future biomarker development and precision medicine approaches in CRC.

**Figure 1:**
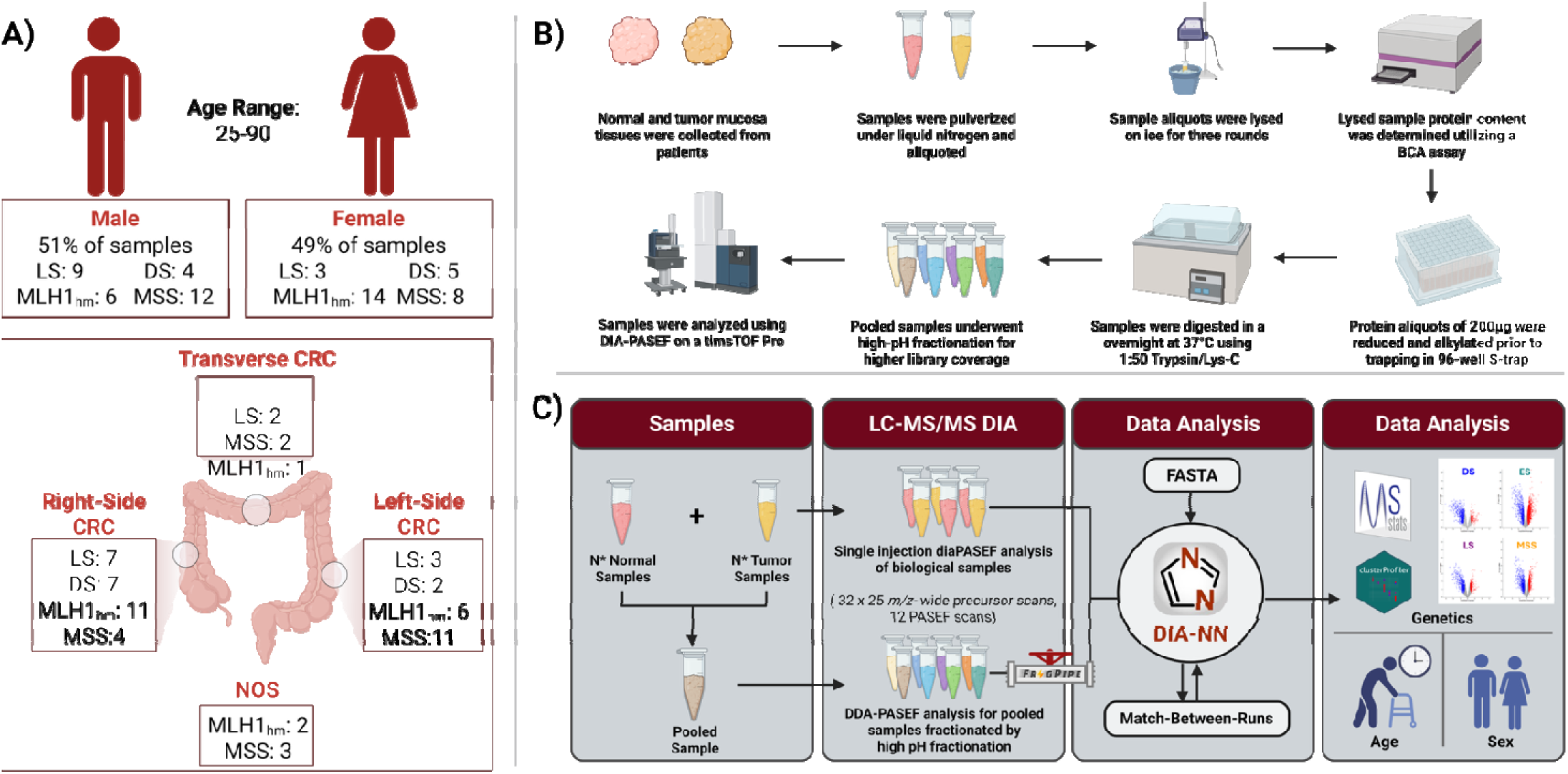
Summary of Samples and Sample Preparation and Analysis. (A) Breakdown of the 122 tumor and normal mucosa samples from the 61-patient cohort across the four colorectal cancer (CRC) subtypes: Microsatellite Stable (MSS), MLH1 Hypermethylation (MLH1_hm_), Lynch Syndrome (LS), and Double Somatic (DS). Additional information on patient age at diagnosis, gender and original tumor location are also provided. **(B)** Overview of sample preparation for patient tumor and mucosa samples. **(C)** Overview of DIA analysis from generation of the pooled samples for mucosa and tumors to data analysis.

## Experimental Procedures

### Tissue Collection and Pulverization

Institutional review board approval was obtained from the Ohio State University (Study #2018C0146). CRC and normal tissues were obtained from the Biospecimen Services Shared Resource of the Ohio State University from participants who consented to the biorepository arm of the Ohio Colorectal Cancer Prevention Initiative (OCCPI). ^14^ The human subjects abide by the Declaration of Helsinki principles. Methods for the OCCPI are described elsewhere, but briefly, 3310 patients who had surgical resection in Ohio for newly diagnosed colorectal adenocarcinoma were enrolled in the OCCPI between 2013 and 2016. A blood and two FFPE tissue samples (tumor and normal mucosa) were obtained from every participant. Fresh frozen tumor and normal tissue was obtained on a subset of patients who were diagnosed at Ohio State University. All patients received MMR deficiency screening and select patients underwent germline genetic testing (including all with MMR deficiency). From the 3310 patients enrolled, 142 were diagnosed with Lynch syndrome. ^5, 14^

We conducted a label-free bottom-up proteomics analysis of 122 fresh frozen tumor and complimentary normal mucosa tissues from three types of MSI CRC samples: LS (n = 12), DS (n = 9), and *MLH1*_hm_ (n = 20), as well as MSS samples (n = 20), from a 61-patient cohort. (**Figure 1 and Table 1**). The entire sample set was blinded and randomized before any further sample preparation was conducted. Tissue chunks were immersed in liquid nitrogen and were pulverized into a fine powder using the Cellcrusher device (Cellcrusher, Schull, Cork, Ireland), which was pre-cooled in liquid nitrogen prior to use. The pulverized solid powder was then transferred to an ultracentrifuge tube, which was kept cooled in dry ice for the entire process. Twenty-four samples were partially un-blinded and grouped as A, B, C or D, which represented MSS, LS, DS or *MLH1*_hm_, respectively, and was designated as Batch 1. Batch 1 was used to generate a pooled sample for high pH Fractionation to create a spectral library. The entire sample set can be found in **Table 1** and **Figure 1**.

### Tissue Lysis and Protein Digestion

Tissue lysis was conducted using a probe sonicator (Fisher Scientific) in pulse mode (1 second on, 2 seconds off) at 20% amplitude for a total of 1 minute sonication per sample. A bicinchoninic acid (BCA) assay was conducted using the Pierce BCA Protein Assay Kit (ThermoFisher Scientific, Rockford, Illinois, USA) to quantify the protein concentration. Afterward, 200 µg of protein for each sample was aliquoted on separate tubes and spun in a vacuum centrifuge to dryness. The protein digestion steps were conducted using the Suspension-Trapping method (ProtiFi, Fairpoint, New York, USA) with a few modifications.^15^ Briefly, each sample was resuspended in 50 µL lysis buffer containing 5% SDS and 5 mM triethylammonium bicarbonate (TEAB). DTT was added with a final concentration of 20 mM and samples were incubated in a thermomixer at 95 °C for 10 minutes. Iodoacetamide was added with a final concentration of 40 mM and samples were incubated at room temperature for 30 minutes in the dark and then centrifuged for 8 minutes at 13,000 x g. Afterwards, 5 µL of 12% phosphoric acid was added to each sample and vortexed for a few seconds before adding 350 µL of a buffer solution containing (9:1) methanol and 1M TEAB. The resulting solutions were transferred individually to a corresponding well in the 96-well S-Trap digestion plate. The plate was subjected to a 1,500 g for 2 minutes in a centrifuge. The proteins were trapped in the filter of the plate and were washed with 200 μL of binding buffer, followed by centrifugation at 1,500 g for 2 minutes for each cycle. Afterward, the S-Trap digestion plate was placed on top of a clean receiver plate and 125 μL of digestion buffer, which contained 1:50 wt: wt a mix of Trypsin and Lys-C protease mix. The plate was incubated overnight at 37 °C in a humid environment. After the incubation, an additional 80 μL digestion buffer was added to all wells of the S-Trap digestion plate and was centrifuged at 1,500 g for 2 minutes. The eluted peptides in the collection plate were then subjected to vacuum centrifugation to dryness and stored at -20 °C until LC-MS/MS analysis.

### High-pH Fractionation of the Pooled Sample Group

To increase the protein sequence coverage of the study, protein digests from Batch 1 were subjected to an offline high-pH reversed-phase liquid-chromatography (RPLC) fractionation using the Pierce™ High pH Reversed-Phase Peptide Fractionation Kit (ThermoFisher Scientific, Rockford, Illinois, USA). A pooled digest was created by taking an equal volume from each sample in the Batch after resuspending in 0.1% formic acid. Following the manufacturer’s protocol, the sample was loaded into the fractionation spin column and eluted by different percentages of acetonitrile/water solution, resulting in 8 peptide fractions. This enabled the generation of a set of peptide fractions representing an equal representation of different patient cohorts analyzed by RPLC-MS/MS.

### RPLC-MS/MS Analysis

Prior to RPLC-MS/MS analysis, samples were resuspended in 0.1% formic acid. Protein identification was performed using nanoflow RPLC-MS/MS on a Bruker timsTOF Pro equipped with a CaptiveSpray source operated in positive ion mode. Samples were separated on a C18 RPLC column (1.6 µm particle size, 75 µm i.d. / 360 µm o.d., 250 mm length; IonOpticks) using a Bruker nanoElute UHPLC system. The column was equilibrated with 4 column volumes at 800 bar prior to each sample injection. Mobile phase A was 0.1% formic acid in water; mobile phase B was 0.1% formic acid in acetonitrile. A flow rate of 0.4 µL/min was used. Mobile phase B was increased from 2% to 17% over the first 40 minutes, to 25% over the next 20 minutes (60 min total), to 37% over the next 10 minutes (70 min total), and to 80% over the next 10 minutes (80 min total), then held at 80% for a final 10 minutes, for a total 90-minute gradient. MS and MS/MS spectra were recorded over m/z 100–1700 and 1/K0 0.6–1.6. Data-independent acquisition (DIA) with parallel accumulation-serial fragmentation (PASEF)^16, 17^ was used for data acquisition, with 32 mass steps per cycle over the range 400–1200 m/z and a mobility range of 0.61–1.49 1/K0, using a mass width of 25 Da with no mass or mobility overlap. Collision energy was ramped with increasing 1/K0, from 20 eV at 0.60 1/K0 to 59 eV at 1.60 1/K0. Given the high sample number and the 90-minute gradient length, each sample was analyzed once. One patient’s matched normal mucosa sample failed to yield a usable signal at acquisition and was excluded prior to downstream processing; the corresponding tumor sample was retained through spectral library searching but was subsequently excluded from differential abundance analysis due to the lack of a matched normal. The final paired dataset comprised 122 mass spectrometry runs from 61 patients (**Table 1).**

### Spectral Library Generation using FragPipe and Identification and Quantification in DIA- NN

Study-specific spectral library was generated from data-dependent acquisition (DDA) MS/MS data using the FragPipe computational platform (version 18.0).^18^ High-pH reverse-phase fractionated samples were used, comprising 8 fractions from Batch 1 acquired in triplicate (24 total DDA acquisitions). Protein identification was performed using nano-liquid chromatography-nanospray tandem mass spectrometry (LC/MS/MS) on a Bruker tims-ToF Pro equipped with a CaptiveSpray source operated in positive ion mode. Samples were separated on a C18 reverse phase column (1.6 µm, 250mm* 75 µm IonOpticks) using a Bruker nanoElute UHPLC system. Pre-injection the column was equilibrated with 4 column volumes at 800 bar. Mobile phase A was 0.1% Formic Acid in water and acetonitrile (with 0.1% formic acid) was used as mobile phase B. A flow rate of 0.4 µL/min was used. Mobile phase B was increased from 2 to 17% over the first 40 minutes, then increased to 25% over the next 20 minutes, further increased to 37 % over the next 10 minutes, and finally increased to 80 % over 10 minutes and then held at 80 % for 10 minutes.

MS and MS/MS experiments were recorded over the m/z range 100-1700 and 1/K0 of 0.6-1.6. DDA-PASEF was used for all experiments, the overall acquisition cycle was 1.16 seconds and comprised of a tims-MS scan and 10 PASEF MS/MS ramps, a polygon filter was applied to the m/z and IM dimensions to exclude singly charged species. Active exclusion was applied, releasing after 0.4 minutes, with precursor reconsidered if current intensity/previous intensity was 4.0 or greater. Collision energy was increased with increasing 1/K0 and was set to 20 e1/K0 at 0.60 increasing to 59 at 1.60 1/K0.

Raw DDA files were searched using FragPipe (MSFragger search engine) against a curated UniProt human reference proteome database (accessed March 2022; 147,520 protein entries), appended with common contaminants and reversed-sequence decoys (prefix "rev_"). Common internal Retention Time standards (CiRT) alignment was enabled during library generation to align retention times across fractionated runs, including technical replicates. A target-decoy approach was used to filter peptide and protein identifications.^19^ Search parameters specified strict trypsin digestion (maximum 2 missed cleavages), peptide lengths of 7–50 amino acids, and a precursor mass range of 500.0–5000.0 Da. Carbamidomethylation of cysteine (+57.0215 Da) was set as a fixed modification; variable modifications included oxidation of methionine (max. 3), protein N-terminal acetylation, and pyroglutamate formation. Following automated internal mass calibration, search tolerances were set to 20 ppm (precursor) and 10 ppm (fragment), scoring b- and y-ion series. The library was filtered to 1% FDR at both the protein and peptide levels, yielding a spectral library of 7,744 proteins, which was imported into DIA- NN.^20^

The 123 experimental DIA files — comprising 122 paired tumor-normal runs from 61 patients plus one unpaired tumor sample whose matched normal failed acquisition — were analyzed in DIA-NN (version 1.8.1) against this study-specific spectral library, which was re-annotated against the human UniProt FASTA database to ensure accurate site localization of terminal modifications. Precursor detection and peak selection were performed using DIA-NN’s integrated deep neural networks, with mass accuracy, retention-time alignment, and scan-extraction windows dynamically inferred and optimized per run to account for instrument calibration drift. N-terminal acetylation was actively monitored for site localization. A strict 1% (0.01) FDR threshold was applied globally at the precursor and protein levels using a target-decoy competition strategy, with highly heuristic protein grouping based on protein names (pg-level 1) to reduce the number of protein groups obtained. Full search parameters are provided in the **Supplemental Information, Table S3**.

### Bioinformatics Analysis: Differential Analysis in MSStats and Perseus

The DIA-NN output matrix — comprising 7,352 proteins identified and quantified across the full set of tumor and matched normal mucosa samples, out of the 7,744-protein spectral library — was analyzed in the R-based MSstats package (version 4.0) together with an accompanying annotation file specifying tissue condition ("tumor" or "normal") and CRC subtype for each sample. MSstats was run using a "paired" experimental design, reflecting that each subject contributed both a tumor and a matched normal biopsy; the one unpaired tumor sample lacking a matched normal was excluded at this stage, yielding a final dataset of 122 paired runs from 61 patients. Raw precursor and fragment ion intensities from DIA-NN were imported using MSstats’ DIA-NN converter with Match Between Runs (MBR) enabled, retaining only features with Global, Library, and Library Protein Group Q-values < 0.01. Data filtering removed proteins identified by a single feature only and features with fewer than three measurements across all runs; where a feature had multiple measurements within a single run, the maximum intensity was retained. Missing-value imputation was performed using the accelerated failure time (AFT) model implemented in MSstats v4.0,^21^ in which log-transformed intensities below a noise cutoff (q□₀.₁ = 1.0116 in this dataset) were treated as censored missing values; this cutoff is automatically estimated by MSstats from the 25th, 75th, and 99.9th percentiles of each dataset’s own log_2_-intensity distribution (q□₀.₁ = q□₂₅ − (q□₉₉.₉ − q□₇₅)), rather than manually specified.^21^ Equalize-medians normalization was applied to correct for systematic technical variation across MS runs, including batch-to-batch (day-to-day) acquisition differences. Protein-level abundance summaries were calculated using Tukey’s Median Polish (TMP), using the top three most intense features per protein. Statistical significance of differential protein abundance between tumor and normal conditions was assessed using linear mixed-effects models, accounting for biological variance between patients (modeled as a random effect nested within condition) and residual technical variance between runs; because tumor and matched normal samples were always acquired within the same batch for a given patient, batch was fully confounded with subject in this design and was not modeled as an independent variance component. Resulting p-values were adjusted for multiple testing using the Benjamini-Hochberg procedure, with significance defined at an adjusted p-value < 0.05. Protein-level abundances and the full MSStats differential abundance results are available in **Supplemental Information, Table S4**.

Statistically significant proteins for each subtype were filtered and visualized their overlap using Venny 2.1, ^22^ where we obtained the 260 proteins list which we considered the shared dysregulated proteins between the CRC subtypes. We further investigated the differential protein expressions of these proteins by using the Abundance Profiles feature in Persesus. ^23^

### GO/KEGG analysis

In the analysis conducted using the ’clusterProfiler’ R package ^24^, the ’enrichKEGG’ function was employed with ’hsa’ specified as the organism (representing Homo sapiens) and ’org.Hs.eg.db’ selected as the organism database. To adjust p-values for controlling the FDR, the Benjamini-Hochberg (B.H.) procedure was utilized. The analysis set a p-value cutoff of 0.05 to identify statistically significant pathways, alongside a q-value cutoff of 0.2 to further mitigate the risk of false positives in the context of multiple hypothesis testing.

Similarly, the ’enrichGO’ function from the same package was utilized, with ’org.Hs.eg. db’ serving as the organism database. The B.H. method was again chosen for FDR adjustment. This function was applied to all three main Gene Ontology (G.O.) categories—Biological Process, Molecular Function, and Cellular Component—under the collective designation "ALL". The criteria for statistical significance were maintained with a p-value cutoff of 0.05 and a q-value cutoff of 0.2, ensuring a consistent approach to controlling false discoveries across both enrichment analyses.

### Graph analysis

To better understand the functional pathways underlying protein dysregulation in each CRC subtype, Gene Ontology (GO) enrichment analyses were performed on two tiers of protein sets: (1) significantly dysregulated proteins within each subtype, and (2) the top upregulated and downregulated proteins shared across all subtypes.

We next examined the statistical significance of sex-based differences in protein dysregulation across the CRC subtypes. P-values were calculated to evaluate how protein dysregulation levels differed between male and female patients, with lower p-values (-log_10_-transformed) indicating greater sex-associated differences. The analysis focused on the top proteins most significantly associated with sex-based dysregulation, visualized in the heatmap (**Figure 6**) and selected from a larger dataset, which included LS = 69, DS = 19, MLH1_hm_ = 63, and MSS = 10 proteins.

### Analysis of Ribosomal Protein Expression

This analysis utilizes results from KEGG/GO analyses to identify proteins associated with ribosomes across various subtypes. A clustered heatmap is generated to visualize the abundance of these ribosomal-related proteins, employing an "Average" clustering method. This method calculates the distance between two clusters as the mean of all distances between their elements, enabling the identification of distinct expression patterns and similarities in ribosomal protein abundance across subtypes.

### PCA Plot

For each subtype, the initial step involves identifying and matching each protein to its corresponding gene, while discarding proteins that lack gene matches. Subsequently, leveraging results from GO/KEGG analysis, the log2Fold Change (log2FC) data for each protein is compiled and cross-referenced with the Log (Intensity) information. The matched proteins are then utilized for Principal Component Analysis (PCA) plotting. The PCA plot creation process begins with standardizing the data, followed by reducing its dimensionality to two principal components. These components are visualized in a scatter plot, effectively illustrating the data’s distribution and variance within a two-dimensional space, providing a comprehensive view of the underlying biological processes.

### Age-Related Expression Analysis

For each CRC molecular subtype, patient-specific protein dysregulation was calculated as the normalized, log_2_-transformed protein abundance in the tumor minus that in the corresponding matched normal mucosa from the same patient. Age-correlation analysis was performed within each subtype’s set of proteins significantly dysregulated between tumor and matched normal tissue (LS: n = 648; DS: n = 1,036; MLH1_hm_: n = 906; MSS: n = 604 significant proteins), restricting the analysis to proteins already established as robustly tumor-associated within that subtype. Within each subtype-specific set, Pearson correlation was calculated between age at diagnosis and patient-specific tumor-normal dysregulation for every protein. Given the modest per-subtype cohort sizes (n = 9–20 patients; Table 1), we selected an effect-size threshold (|r| ≥ 0.5) rather than a nominal p-value cutoff to identify age-associated proteins, since p-values from Pearson correlation are highly sensitive to sample size and can be underpowered to detect even moderate-to-strong associations at these cohort sizes. Proteins meeting |r| ≥ 0.5 were retained (LS: n = 133; DS: n = 134; MLH1_hm_: n = 42; MSS: n = 35) and ranked by absolute correlation coefficient; the 10 proteins with the strongest |r| per subtype were selected for visualization in a merged heatmap (**Figure 4**).

### Sex-Specific Expression Analysis

For each patient and protein, protein dysregulation was calculated as the normalized, log_2_-transformed protein abundance in the tumor minus that in the corresponding matched normal tissue. Sex-associated analysis was performed within each subtype’s set of proteins significantly dysregulated between tumor and matched normal tissue (LS: 648 proteins; DS: 1,036; MLH1_hm_: 906; MSS: 604), restricting the analysis to proteins already established as robustly tumor-associated within that subtype. Within each subtype-specific set, male and female patient-specific dysregulation values were compared using an independent two-sample Student’s t-test, with p < 0.05 as the significance threshold. Proteins meeting this criterion were ranked by p-value in ascending order, and the 10 proteins with the smallest p-values per subtype were selected for visualization (LS = 21, DS = 24, MLH1_hm_ = 15, and MSS = 15 proteins met this threshold in total). Heatmaps display −log_10_-transformed p-values to illustrate the strength of sex-associated differences in protein dysregulation (Figure 6); a combined heatmap integrating the selected proteins across all four subtypes was also generated. The complete dataset was deposited to the ProteomeXchange Consortium ^25^ via the PRIDE ^26^ partner repository with the dataset identifier PXD073693.

### Gene Expression Analysis

To evaluate gene expressions in colorectal cancer comparatively to the reported dataset, the TCGA-COAD data was loaded into UCSC Xena Database for analysis. ^27^ Briefly, the TCGA Colon Cancer dataset (n=551) was chosen as the first dataset with a first variable of phenotypic response of ‘sample_type’. Then, each gene of interest in the top ten up- and down-regulated genes were added individually to the database with ‘gene expression’ analysis given. Then, all “null” data points were removed from the gene expression leaving n=329 samples for analysis through the four subtypes of recurrent tumor, primary tumor, metastatic, and normal tissue. Gene expressions for each protein were exported for analysis into GraphPad Prism 11.1.0 and data was filtered to only look at statistical significance between normal and primary tumor tissues (n=327) to determine if genes were up or down regulated within tumor tissues as compared to normal using an unpaired t-test. These datasets can be found in the **Supplemental Information, Table S5**.

### Protein Data Consortium (PDC) Protein Expression Analyses

To evaluate previously obtained clinical data to aid in validation of proteomics data within the manuscript, the web-based platform Cancer Proteogenomic Data Analysis Site (cProsite) ^28^ was used to obtain protein expression levels from the CPTAC-2 dataset for colon cancer. Briefly, relative protein abundance was obtained for proteins of interest and data for each sample was exported to GraphPad Prism 11.1.0. For sex-associated analyses, patient ID values were utilized to correlate demographic information collected at the time of the study and non-recorded values were not utilized in the analysis. An unpaired t-test was utilized to evaluate patient samples (n=101) tumor versus adjacent normal protein abundances. In the case of sex-associated analyses, these were separated by grouped columns. This dataset can be found in Supporting Figure 10.

## RESULTS

The samples were acquired through The Ohio Colorectal Cancer Prevention Initiative. ^14^ The CRC samples included in this study are novel due to their geographical specificity and contain a diverse age group. This cohort included rich metadata to better understand the phenotypic and proteomic data shared in this study. We uncovered dysregulated proteins across all four CRC subtypes, with subtype-specific expression patterns that highlight potential diagnostic biomarkers. As expected, our findings reveal substantial differences between the proteomes of MSS and MSI-high patients. We also observe age-associated shifts in metabolic and signal transduction pathways, particularly in the DS subtype.

### Global protein expression differences between CRC subtypes

Analysis of whole proteome expression alterations between tumor and normal tissues revealed 1,607 significantly dysregulated proteins. Dysregulated proteins are defined as proteins with a log_2_ fold-change greater or less than 1 and resulting p-values were adjusted for multiple testing using the Benjamini-Hochberg procedure, defining significance at an adjusted q-value < 0.05 between tumor and corresponding normal mucosa, in at least one CRC subtype. Within each subtype, we identified 1,036, 648, 906, and 604 dysregulated proteins for the Double Somatic (DS), Lynch Syndrome (LS), MLH1 hypermethylation (MLH1_hm_), and Microsatellite Stable (MSS) subtypes, respectively (**Figure S1**). Among these, 374, 69, 203, and 134 proteins, respectively, were uniquely dysregulated in each subtype, while 260 dysregulated proteins were common to all subtypes (**Figure 2A**). Taking a closer look at the 260 shared dysregulated proteins, hierarchical clustering was performed, generating a heatmap that distinctly separates CRC subtypes based on their log_2_ fold-change profiles (**Figure 2B**). The detected protein changes in the MSS cohort were significantly differentiated from the three MSI-high patient cohorts (**Figure 2C**), with branching in the heat map showing MSS as distinct from the three MSI-high variants. Within the three MSI-high cohorts, LS and MLH1_hm_ were more similar, compared to the DS group. A complete list of all dysregulated proteins is available in the **Supplemental Information, Table S1**. **Supplemental Figures S4 – 6** show the Top Biological Processes from Gene Ontology analysis in MSS, MLH1_hm_, and DS subtypes, respectively.

**Figure 2:**
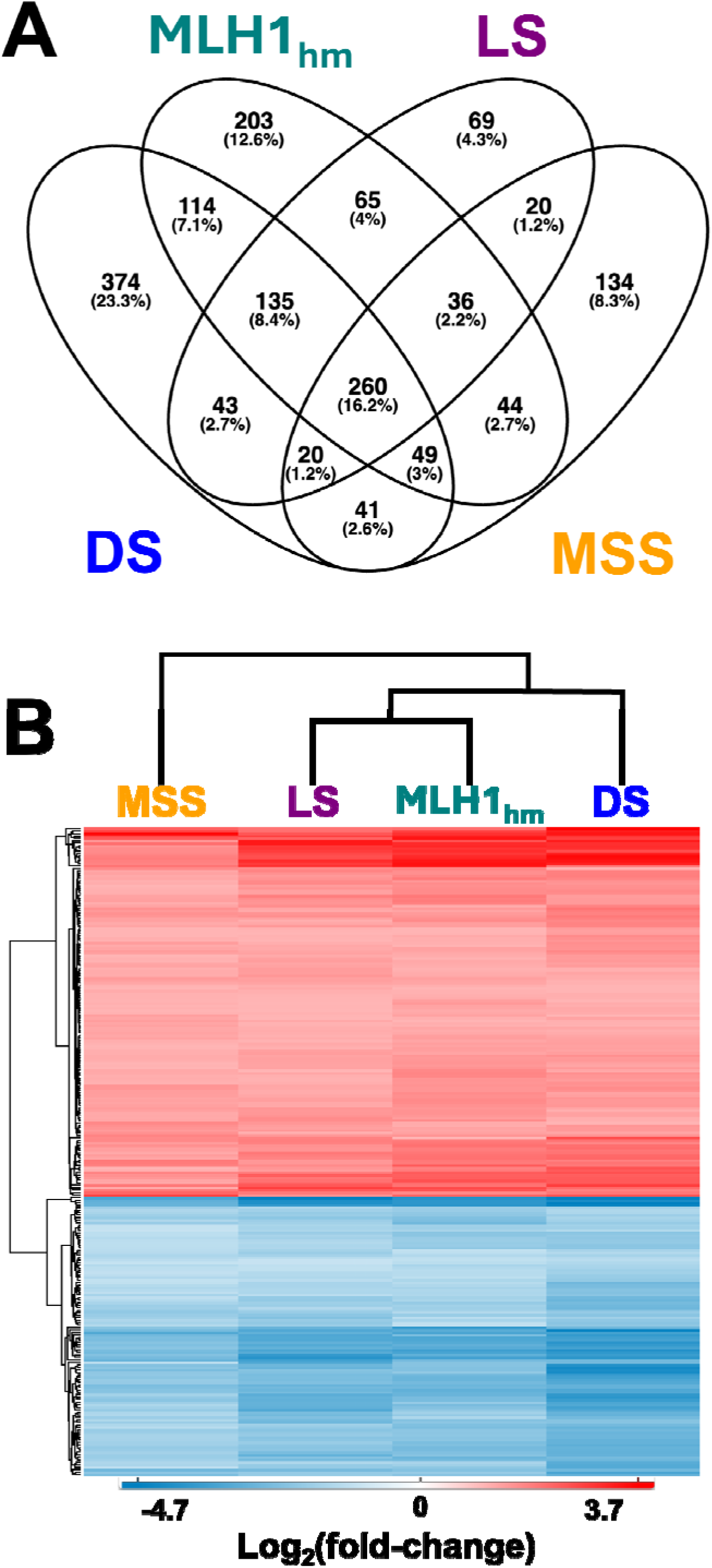
Overlap and clustering of dysregulated proteins across CRC subtypes. (A) Venn diagram showing the overlap of the 1,607 proteins significantly dysregulated (Benjamini-Hochberg adjusted p-value < 0.05 and |log₂FC| > 1, tumor vs. matched normal mucosa) in at least one of the four colorectal cancer (CRC) subtypes: Double Somatic (DS; 1,036 proteins), MLH1 Hypermethylation (MLH1_hm_; 906), Lynch Syndrome (LS; 648), and Microsatellite Stable (MSS; 604). Each region shows the number of proteins unique to or shared between subtypes, with percentages relative to the total of 1,607. The central intersection contains the 260 proteins dysregulated in all four subtypes. (B) Hierarchical clustering heatmap of these 260 shared proteins based on their log_₂_ fold-change (tumor vs. matched normal) in each subtype. Red indicates upregulation and blue indicates downregulation in tumor relative to matched normal mucosa (scale: −4.7 to 3.7). The column dendrogram separates MSS from the three mismatch repair-deficient subtypes (LS, MLH1_hm_, and DS), with LS and MLH1_hm_ clustering most closely.

Lynch Syndrome is the predominant form of hereditary colorectal cancer and increases susceptibility to a range of secondary malignancies. Identifying LS status is essential for guiding targeted cancer therapies and implementing preventative screening for both patients and their families. Of the 7,352 proteins identified and quantified across the full tumor and normal mucosa sample set, the large majority were quantified with a valid tumor-matched normal comparison within each individual CRC subtype: 7,232 proteins in LS, 7,175 in DS, 7,275 in MLH1_hm_, and 7,292 in MSS. The remainder were either detected in only one of the two tissue types within a given subtype (114, 163, 73, and 58 proteins in LS, DS, MLH1_hm_, and MSS, respectively, precluding a finite fold-change estimate) or were not detected at all in that subtype (6, 14, 4, and 2 proteins, respectively). To identify the prominent protein alterations characteristic of the LS subtype, we evaluated the top 10 most upregulated and downregulated proteins based on a strict threshold (log_2_FC ≥ |1|, adj. p-value ≤ 0.05) as shown in **Table 2**. The LS subtype demonstrates a distinct expression profile led by profound downregulations in extracellular matrix and metabolic regulators such as OGN (log_2_FC = -4.58, adj. p-value = 2.45E-07) and PGM5 (log_2_FC = -3.94, adj. p-value = 1.19E-06), alongside prominent upregulations in proteins including RPL22L1 (log_2_FC = 3.47, adj. p-value = 1.37E-05). When comparing this signature across the other subtypes, we observed that most of these top-altered proteins are shared globally across the subtypes, pointing toward conserved downstream pathophysiological mechanisms. For instance, extracellular matrix markers PRELP, OGN, and PGM5 remain heavily downregulated across all three other subtypes, with DS showing the largest magnitude of downregulation for OGN (log₂FC = −4.73) and PRELP (log₂FC = −4.53). Their relative log₂FC values in LS are modest (−0.29 to −0.66) compared with their large absolute fold-changes, and several other top proteins, including AOC3, GNAO1, and AZU1, have relative values near zero (+0.02 to +0.03), indicating that their dysregulation in LS is potentially characteristic of CRC broadly. However, a few notable exceptions highlight subtype-specific molecular differences. HKDC1 is significantly upregulated in LS (log_2_FC = 2.72, adj. p-value = 0.038), but loses statistical significance in MSS and MLH1_hm_ groups, where its expression shifts (MSS log_2_FC = -1.91, adj. p-value = 0.203; MLH1_hm_ log_2_FC = 1.70, adj. p-value = 0.062). As mentioned prior, RPL22L1 exhibits high significance in LS, but also in MLH1_hm_ (log_2_FC = 3.00, adj. p-value = 2.42E-06), while remaining statistically non-significant in the MSS subtype (log_2_FC = 0.31, adj. p-value = 0.708). These shared patterns alongside sub-type specific protein alterations underscore both the unified proteomics landscape of these colorectal cancers and the finer differences distinguishing LS tumors from sporadic subtypes.

Data from proteomics analyses were compared to data obtained in the cancer genome atlas colon adenocarcinoma (TCGA-COAD) gene expression analyses. ^29^ For the top ten reported down-regulated proteins, all ten genes showed the same expressions in the TCGA-COAD with statistical significance. Comparatively, among the top ten up-regulated proteins observed, only six matched in expression within this dataset. Furthermore, of the four datasets that did not match in expression, gene PRTN3 was not observed as significant within the gene expression cohort. Plotted gene expressions from the TCGA-COAD analysis of primary tumors and normal solid tissues can be found in **Supplemental Figures 2-3**. Comparatively, when pulling proteomics expression analyses from CPTAC-2 within the National Cancer Institute Proteomic Data Commons (PDC), ^29^ all protein expressions that are observed within the LS cohort exhibit similar trends of up and down regulation (**Supplemental Figures 8-9**).

We next identified protein alterations unique to the LS subtype to distinguish proteomic alterations specific to germline MMR-deficient subtype from changes across MSS, double-somatic MMR-deficient and MLH1_hm_ epigenetically silenced subtypes. We evaluated the statistically significant proteins based on the same strict threshold of log_2_FC ≥ |1|, adj. p-value ≤ 0.05 and found 69 proteins (51 upregulated, 18 downregulated) altered between tumor and normal mucosa from LS patients. Among the upregulated proteins, several functional groups stood out beyond the pan-CRC proliferative signature already noted: immune/inflammatory regulators not shared with the other subtypes (LRBA, APOL2, CASP1, BTK, TRIM22, STAT2), additional cell-cycle and mitotic proteins (TOP2A, KIFC1, KIF2A, RFC4, NCAPH/condensin, LARP1), RNA-processing and splicing factors (PRPF38A, SNRPA, HNRNPA0, FAM133B), molecular chaperones (DNAJB6, DNAJC9, PPID), and metabolic enzymes tied to glycolysis and nucleotide salvage (LDHA, PFKFB3, HKDC1, ADI1, UCK2). Notably, cathepsin K (CTSK) and MMP12 point to a distinct matrix-degrading enzyme profile, while DACH1 and CIC, which are both transcriptional regulators with tumor-suppressive roles in other cancers, were also uniquely elevated.

The smaller set of LS-unique downregulated proteins included endothelial/coagulation-associated factors (PROCR, PLAT), additional ECM/basement-membrane components not captured in the pan-subtype signature (COL5A2, HAPLN1, LTBP4), the tight-junction protein claudin-7 (CLDN7), the dendritic-cell marker CD209, and cytoskeletal proteins (TUBA1A, MYO1A, AHNAK2), alongside PYGM (muscle glycogen metabolism) and the bicarbonate transporter SLC4A4. Together, these 69 proteins represent a candidate LS-specific proteomic signature layered on top of the broader tumor-versus-normal mucosa changes and form the basis for the downstream GO enrichment analysis.

The proteins directly substantiate the biological pathways governing this subtype, specifically highlighting a coordinated shift toward metabolic rewiring, translational hyperactivation, immune priming, and extracellular matrix (ECM) remodeling. Key glycolytic and metabolic enzymes within this list, including LDHA, PFKFB3, and HKDC1, function as the core molecular engines driving aerobic glycolysis (the Warburg effect), enabling LS tumors to rapidly generate the ATP and macromolecular precursors required for continuous cellular expansion. ^30^ To meet the immense biomass demands of this hyper-proliferative state, the unique proteome features a robust cluster of ribosomal assembly and nucleolar regulatory proteins (such as RPL30, RPL38, IMP4, and LARP1) alongside essential DNA replication and chromosomal segregation factors like TOP2A, RFC4, and NCAPH. ^31^ Furthermore, the distinct presence of ECM-modifying enzymes (MMP12, CTSK, TIMP2) and structural components (COL5A2) indicates active stromal degradation and desmoplastic invasion characteristic of germline mismatch repair-deficient tumors (Eide et al., 2021). Finally, the identification of immune and inflammatory mediators like CASP1, STAT2, and TRIM22 provides explicit mechanistic support for the strong immunogenic nature and innate "viral mimicry" response in LS, where severe genomic instability continuously primes localized leukocyte surveillance and antiviral-like signaling cascades. ^32^

### Biological Process Enrichment Analysis of Dysregulated Proteins

To identify highly confident biological processes (BP) associated with the LS subtype, over-representation analysis (ORA) using Gene Ontology (GO) was conducted using the R language package clusterProfiler with stringent statistical thresholds. To account for multiple hypothesis testing, p-values were adjusted using the Benjamini-Hochberg false discovery rate (FDR) procedure. Only terms meeting a strict cutoff of adjusted p < 0.05 and q < 0.2 relative to the human proteome were retained. This rigorous filtering ensured that the top enriched pathways reflected robust, high-confidence biological themes rather than detection artifacts. **Figure 3A** illustrates the enrichment analysis results on all 648 significantly dysregulated proteins in LS tumors relative to matched normal mucosa. This analysis returned 208 significantly enriched BP terms. The four most significant and highest-ranking terms by gene ratio were all related to ribosome/ribonucleoprotein biogenesis: ribonucleoprotein complex biogenesis (72/626 genes, p.adjust = 1.54E-23), ribosome biogenesis (62/626, p.adjust = 1.46E-26), rRNA metabolic process (56/626, p.adjust = 7.91E-26), and rRNA processing (51/626, p.adjust = 2.65E-25) — consistent with the large cluster of nucleolar and pre-ribosomal proteins (DDX10/18/21, BOP1, BYSL, WDR-family, NOP2, NPM1/3, UTP3/4/14A) upregulated in tumor tissue. A second cluster involved extracellular matrix and structural remodeling — extracellular matrix organization, extracellular structure organization, and external encapsulating structure organization (each 38/626, p.adjust = 3.29E-8) — reflecting the coordinated loss of structural collagens alongside gain of collagen-modifying enzymes described above. RNA splicing (38/626, p.adjust = 1.40×10⁻□), viral process (37/626, p.adjust = 2.34×10⁻□), and muscle system process (37/626, p.adjust = 1.62×10⁻□) were also significantly enriched, the latter mirroring the marked downregulation of smooth muscle/contractile proteins (ACTA1, MYH11, MYLK, CNN1, TAGLN) noted previously.

**Figure 3:**
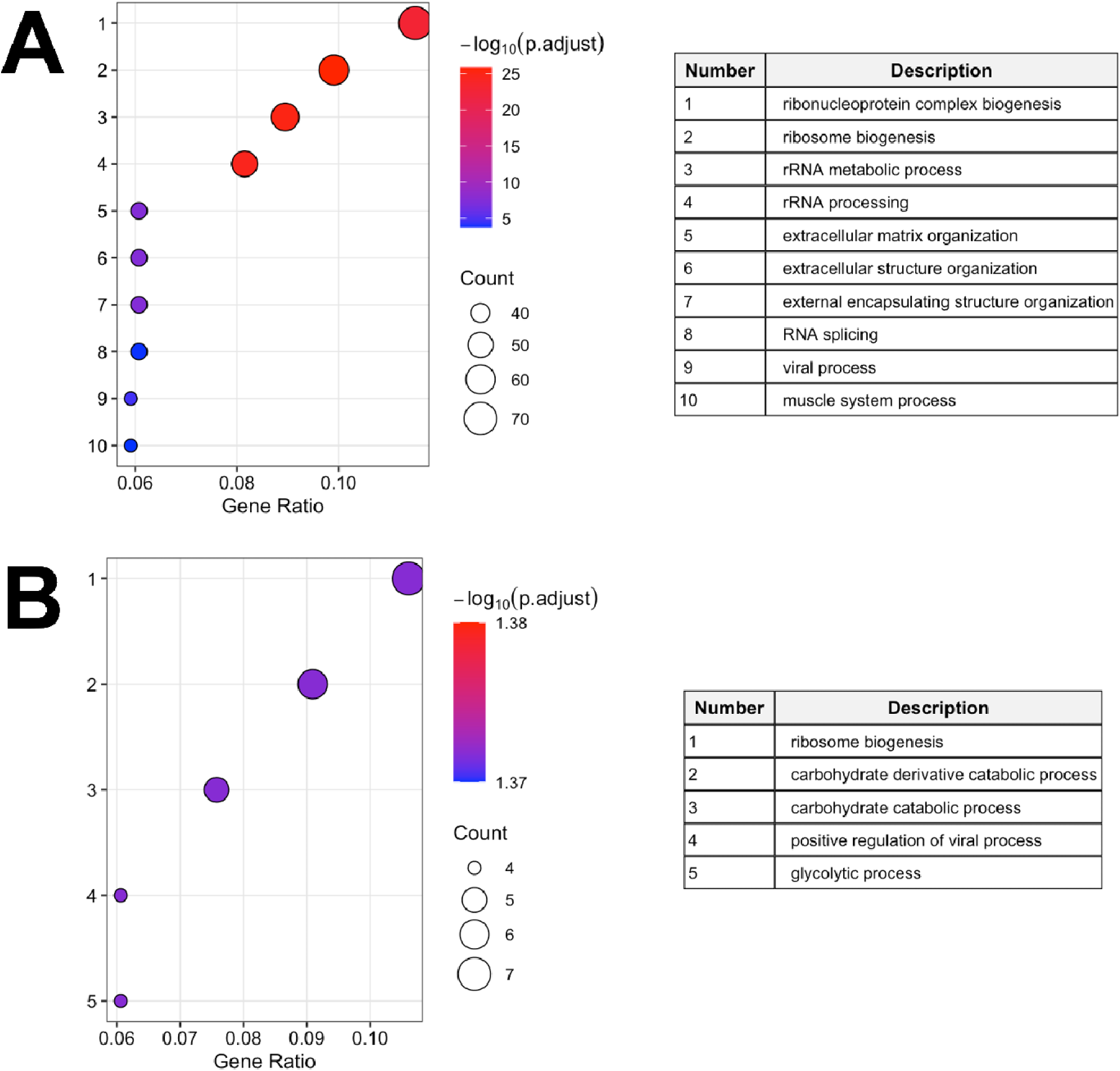
Gene Ontology Biological Process enrichment for the Lynch Syndrome (LS) subtype. Over-representation analysis was performed with clusterProfiler using Benjamini-Hochberg correction (adjusted p-value < 0.05, q-value < 0.2) against the human genome background. Terms are numbered and ranked by gene ratio (the number of input proteins annotated to a term divided by the number of input proteins with a GO annotation), and descriptions are listed in the accompanying tables. Bubble size indicates the number of proteins annotated to each term, and color indicates significance as −log_10_(adjusted p-value). (A) Top 10 of 208 significantly enriched terms among the 648 proteins dysregulated in LS tumors relative to matched normal mucosa (626 with GO annotation). (B) Top 5 of 18 significantly enriched terms among the 69 proteins dysregulated uniquely in LS relative to the MSS, MLH1hm, and DS subtypes (66 with GO annotation). All terms in panel B have similar adjusted p-values (−log_10_ ≈ 1.37–1.38), reflecting the small input list; several terms share overlapping proteins (full results in Supplementary Table [X]).

Next, as shown in **Figure 3B**, a more focused GO analysis was performed on the 69 dysregulated proteins unique to the LS subtype to find unique pathways to this subtype. The analysis returned a much smaller set of 18 significantly enriched terms (top 6 are shown), all with adjusted p-values tightly clustered at the significance boundary (p.adjust = 0.0425–0.0478), reflecting the limited statistical power of a 66-gene input list (three of the 69 proteins did not map to an Entrez ID and were excluded from the test). Despite this, a clear and coherent biological signal emerged. Ribosome biogenesis was again the top term by gene ratio (7/66, p.adjust = 0.0425; AK6, IMP4, NLE1, NMD3, RPL38, SDAD1, WBP11), indicating this process distinguishes LS not only from normal mucosa but from other CRC molecular subtypes as well. The remaining significant terms resolved into two functionally distinct groups rather than 18 independent processes. The majority, 10 of 18 terms, including carbohydrate derivative catabolic process, carbohydrate catabolic process, glycolytic process, ADP catabolic/metabolic process, and several purine/pyridine nucleotide catabolism terms were all driven by the same four genes. These four genes, HKDC1, LDHA, PFKFB3, and SLC4A4 (with MMP12 and BTK additionally contributing) to the broader "carbohydrate derivative catabolic process" term, reflect a single underlying glycolytic/carbohydrate-catabolic signature rather than multiple independent enriched pathways. A second, smaller group of terms (positive regulation of viral process, positive regulation of viral genome replication, regulation of viral life cycle) was driven by CD209, LARP1, PPID, and TOP2A; given that these genes have established roles in immune signaling (CD209), translational control (LARP1), protein folding/mitochondrial permeability (PPID), and DNA topology/replication (TOP2A) rather than antiviral defense per se, this enrichment likely reflects GO annotation of these genes’ documented involvement in viral biology rather than an active antiviral response, and should be interpreted with that caveat. A single additional term, cell cycle DNA replication (DACH1, WIZ, ZPR1), was also significant.

To determine which of these processes distinguish LS, we performed the same analysis for the other three subtypes (**Figures S4–S6**). The four top-ranked LS terms (ribonucleoprotein complex biogenesis, ribosome biogenesis, rRNA metabolic process, and rRNA processing) were also among the top-ranked terms in MSS, MLH1_hm_, and DS, and DNA replication, extracellular matrix organization, and muscle system process were significantly enriched in all four subtypes. Cytoplasmic translation was strongly enriched in MLH1_hm_ (adj. p = 8.7×10⁻²□) and DS (adj. p = 2.8×10⁻¹□), weakly in LS (adj. p = 3.6×10⁻³), and not in MSS. No mismatch repair term was enriched in any subtype; base-excision repair was enriched in DS only, at borderline significance (8 proteins, adj. p = 0.047). Analyses restricted to subtype-unique proteins yielded distinct profiles: the MLH1_hm_-unique set (**Figure S5B)** was dominated by cytoplasmic translation (adj. p = 5.4×10⁻□) and RNA splicing, the DS-unique set (**Figure S6B)** by RNA splicing and processing, transcription elongation, and leukocyte-mediated immunity, and surprisingly, the MSS-unique set returned no significant terms. Glycolytic and carbohydrate catabolic terms appeared only in the LS-unique analysis, supporting this metabolic program as a feature specific to LS.

Taken together, the LS-unique GO signature, once redundant sibling terms are collapsed, reduces to two principal biological themes distinguishing LS from other CRC subtypes: enhanced ribosome biogenesis (shared with the broader tumor-vs-normal signature) and a glycolytic/carbohydrate-catabolic metabolic program (HKDC1, LDHA, PFKFB3, SLC4A4) that is not shared with MSS, double-somatic, or MLH1_hm_ CRC subtypes. The Complete GO Biological Processes results are available in the **Supplementary Information, Table S2.**

### Age-Associated Protein Dysregulation in CRC

In addition to the four different CRC subtypes, this current study also provides a diverse age group of patient samples. To determine whether the magnitude of tumor-specific proteomic dysregulation varies with patient age at diagnosis, we examined the relationship between age and patient-specific tumor-normal protein dysregulation (tumor minus matched normal mucosa, log_2_ scale) among proteins already identified as significantly dysregulated between tumor and normal tissue within each of the four CRC molecular subtypes: Lynch Syndrome (LS; n = 12 patients, ages 25–57, 133 significant proteins), double-somatic (DS; n = 9 patients, ages 27–80, 134 significant proteins), MLH1-hypermethylated (MLH1_hm_; n = 20 patients, ages 58–88, 42 significant proteins), and microsatellite-stable (MSS; n = 20 patients, ages 51–90, 35 significant proteins) (Table 1). Because this dysregulation metric is anchored to each patient’s own normal tissue, correlations with age reflect the extent to which tumor-associated proteomic change scales with age, rather than an age-related shift in baseline protein abundance also present in non-malignant tissue.

Age-correlation analysis was restricted to proteins already identified as significantly dysregulated between tumor and matched normal tissue within each subtype (LS: 648 proteins; DS: 1,036; MLH1_hm_: 203; MSS: 134). Within each subtype-specific set, Pearson correlation coefficients were calculated between age at diagnosis and tumor-normal dysregulation for every protein. Applying an effect-size threshold of |r| ≥ 0.5 identified 133 age-correlated proteins in LS (20.5% of the LS set), 134 in DS (12.9%), 42 in MLH1_hm_ (4.6%), and 35 in MSS (5.8%); the 10 proteins with the strongest |r| from each subtype-specific set are shown in **Figure 4**. The proportion of differentially expressed proteins meeting this threshold ranged from 4.6% to 20.5% across subtypes (LS: 20.5%, DS: 12.9%, MLH1_hm_: 4.6%, MSS: 5.8%), with the two smaller cohorts (LS, n=12; DS, n=9) showing somewhat higher proportions than the two larger cohorts (MLH1_hm_, MSS; n=20 each). This pattern is consistent with reduced sample size increasing the likelihood that a protein’s correlation coefficient exceeds a fixed effect-size threshold by chance and underscores that age-correlated findings in the smaller subtypes should be interpreted with appropriate caution pending validation in larger cohorts.

Each subtype displayed a distinct directional bias among its top age-associated proteins. In LS, the top 10 hits were predominantly negatively correlated with age, led by FCN1 (r = −0.942) and BUD23 (r = −0.870), with HPS3 (r = 0.825) and DACH1 (r = 0.816) as the strongest positive correlates. The DS subtype showed a similar negative skew, led by PPHLN1 (r = −0.921), with DCAF1 as the strongest positive correlate (r = 0.904). The MLH1_hm_ subtype was predominantly negative, led by SEC14L2 (r = −0.792), with RPRD1B as the top positive correlate (r = 0.815). In contrast, the MSS subtype trended positive, most strongly for KIF22 (r = 0.852) and RTKN (r = 0.792), while TCEAL5, CIC, and KIF4A were the strongest negative correlates.

A small number of proteins appeared among the top age-associated hits in more than one subtype. SASH3 was positively correlated with age in LS, DS, and MSS, and TBC1D10C was positively correlated in both MLH1_hm_ and MSS, suggesting a shared age relationship for these proteins independent of molecular subtype. By contrast, PPHLN1 was strongly negatively correlated with age in DS but only weakly positively correlated in MLH1_hm_ and MSS, indicating that for this protein the age relationship is itself subtype-dependent.

### Validation of top age-associated proteins by direct scatter analysis

To visualize the relationships underlying the strongest correlations in Figure 4, we plotted patient-specific tumor-normal protein dysregulation against age at diagnosis for the two proteins with the largest |r| in each subtype, alongside boxplots summarizing the age distribution of contributing samples (**Figure S7**). These plots corroborate the correlations in Figure 4: FCN1 (R² = 0.887) and BUD23 (R² = 0.757) in LS; PPHLN1 (R² = 0.848) and DCAF1 (R² = 0.818) in DS; RPRD1B (R² = 0.665) and SEC14L2 (R² = 0.628) in MLH1_hm_; and KIF22 (R² = 0.726) and RTKN (R² = 0.627) in MSS.

Consistent with the known clinical presentation of each subtype, the age ranges contributing to these correlations differed substantially and showed limited overlap: LS patients were diagnosed at the youngest ages (25–57 years), consistent with earlier-onset presentation in germline MMR deficiency, while MLH1_hm_ patients were diagnosed at the oldest ages (58–88 years), consistent with this subtype’s sporadic, age-associated epigenetic origin; DS (27–80) and MSS (51–90) spanned wider, overlapping intermediate ranges. Given the modest sample sizes, particularly in DS and LS, these age-correlated trends should be interpreted as subtype-specific associations within the observed clinical age range and effect-size-based selection, rather than extrapolated as continuous, generalizable, or independently significance-tested age-disease relationships. ^33^

To characterize the biological processes represented among the age-correlated proteins identified in Figure 4, we performed functional annotation analysis (DAVID; GO Biological Process and KEGG pathways) separately on the positively and negatively age-correlated protein sets (|r| ≥ 0.5) within each subtype, retaining terms with nominal p < 0.05 (**Figure 5).**

**Figure 4:**
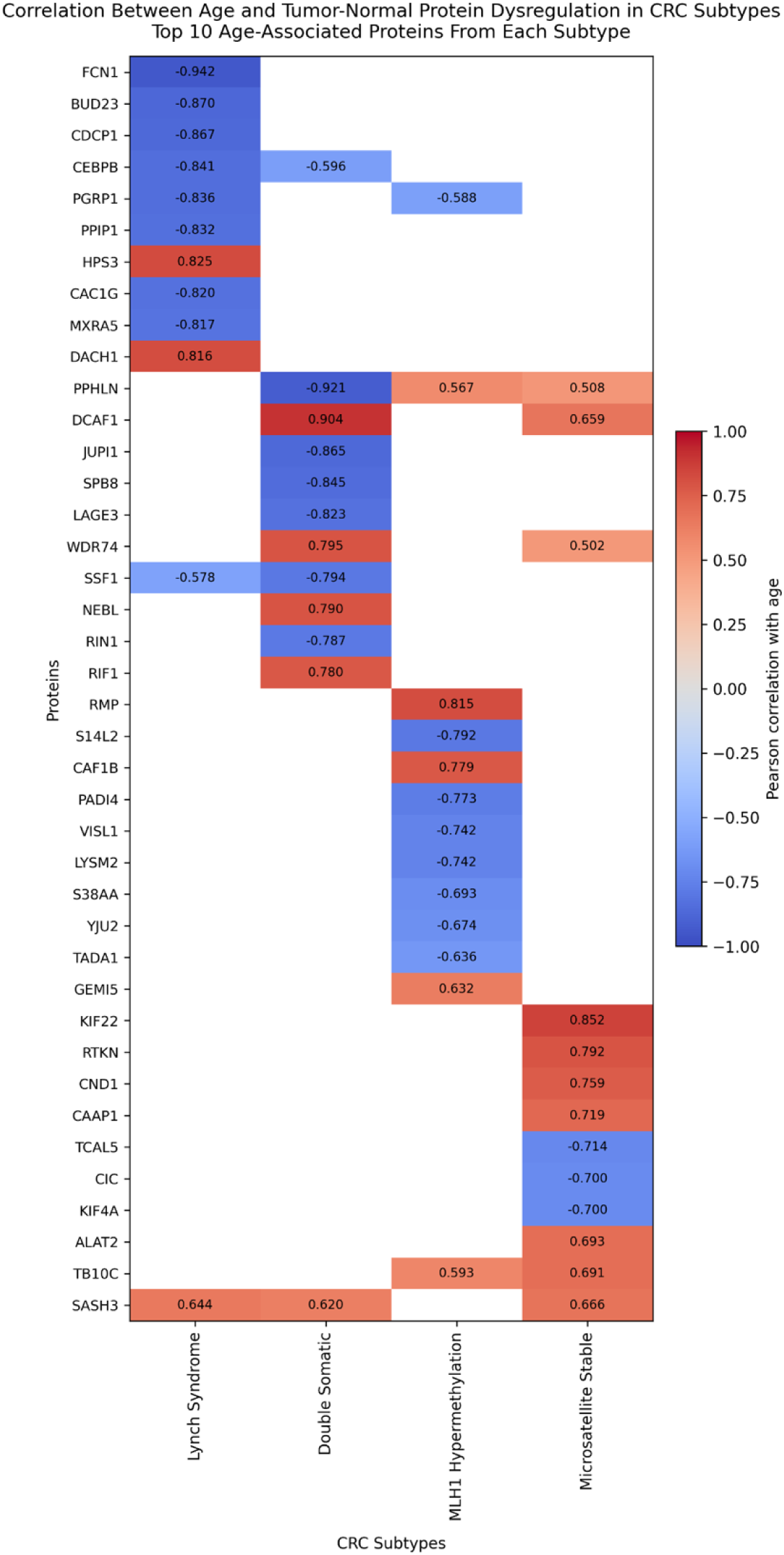
Correlation Between Age at Diagnosis and Protein Expression in CRC Subtypes. This heatmap displays the top 20 proteins most correlated with age at diagnosis in colorectal cancer (CRC) s btypes, highlighting a subset of a larger dataset of dysregulated proteins: Double Somatic (DS) = 141 tota , Lynch Syndrome (LS) = 138, MLH1 Hypermethylation (MLH1_hm_), = 43, and Microsatellite Stable (MSS) = 35. A positive correlation (red) indicates proteins that increase in expression with age at diagnosis, whereas a negative correlation (blue) represents proteins that decrease with age at diagnosis. This visualization highlights age-dependent molecular alterations that may contribute to tumor progression, metabolic shifts, and immune regulation across CRC subtypes.

**Figure 5:**
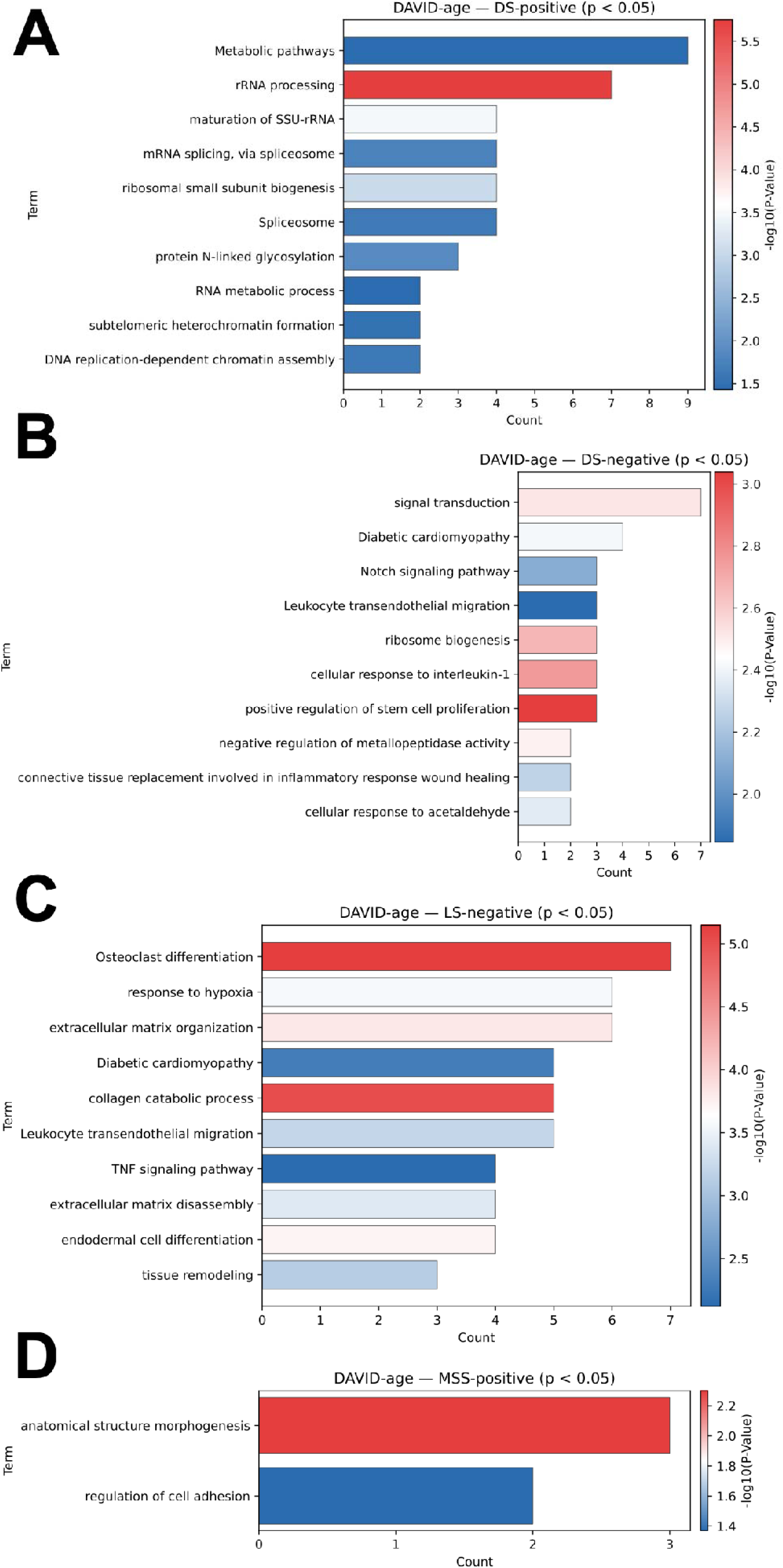
Functional annotation of age-associated proteins. DAVID functional annotation (GO Biological Process and KEGG pathways) of proteins whose tumor-normal dysregulation was correlated with age at diagnosis (Pearson |r| ≥ 0.5), analyzed separately for positively and negatively correlated proteins within each subtype. Bars show the number of proteins annotated to each term, and color indicates significance as −log_10_(p-value); terms with nominal p < 0.05 are shown. (A) Double Somatic (DS), proteins positively correlated with age (n = 83). (B) DS, negatively correlated (n = 51). (C) Lynch Syndrome (LS), negatively correlated (n = 116). (D) Microsatellite Stable (MSS), positively correlated. No significant terms were identified for LS positively correlated proteins, MSS negatively correlated proteins, or MLH1 Hypermethylation (MLH1hm) proteins in either direction. P-values are not corrected for multiple testing; these results are exploratory.

**Figure 6:**
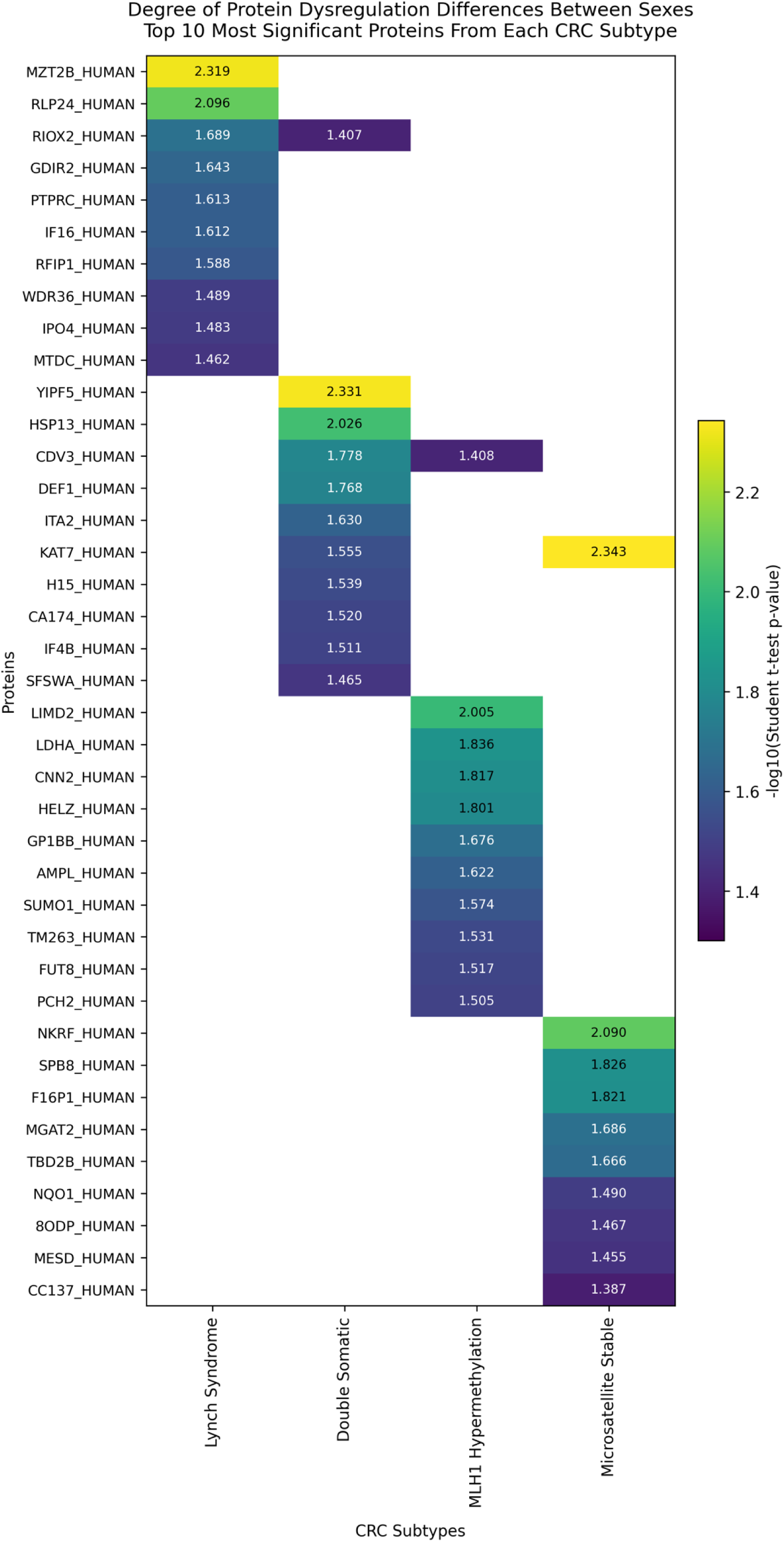
Degree of Protein Dysregulation Differences Between Sexes. This heatmap presents the top proteins most significantly associated with sex-based differences in dysregulation, selected from a larger dataset where DS = 21, LS = 24, MLH1_hm_, = 15, and MSS = 15. The values are represented as -log10(p-values), where higher values (yellow) indicate more significant sex-dependent differences in protein expression, while lower values (purple) suggest minimal differences between male and female patients. This visualization provides insight into sex-associated variability in protein dysregulation, which may be influenced by hormonal, genetic, or immune-related factors that impact CRC tumor biology and potential therapeutic responses.

In the DS subtype, proteins whose tumor-normal dysregulation increased with age (83 of 134 proteins) were enriched for ribosomal and RNA-processing machinery (**Figure 5A**), including rRNA processing (7 proteins, the most significant term in this set), maturation of SSU-rRNA, ribosomal small subunit biogenesis, mRNA splicing via spliceosome, and spliceosome, alongside the broader term metabolic pathways (9 proteins, the largest gene count but weakest significance). This suggests that the proliferative and biosynthetic program characteristic of CRC is more pronounced in older-onset DS tumors. Proteins whose dysregulation instead decreased with age (51 of 134 proteins) were enriched for immune and developmental signaling terms (**Figure 5B**), including signal transduction (7 proteins), Notch signaling pathway, leukocyte transendothelial migration, cellular response to interleukin-1, and positive regulation of stem cell proliferation, indicating relatively greater inflammatory and developmental signaling in younger-onset DS tumors. A smaller ribosome biogenesis term (3 proteins) was also present in this set, indicating that RNA-processing proteins are not exclusively associated with older age in DS.

In LS, significant enrichment was observed only among proteins whose dysregulation decreased with age (116 of 133 proteins; **Figure 5C**); no terms reached significance among the 17 positively correlated proteins. This set was dominated by extracellular matrix remodeling and innate immune processes, including osteoclast differentiation (7 proteins, the most significant term), response to hypoxia, extracellular matrix organization, collagen catabolic process, extracellular matrix disassembly, leukocyte transendothelial migration, TNF signaling pathway, and tissue remodeling. Several of these terms share overlapping proteins, including matrix metalloproteinases (MMP2, MMP8, MMP9, MMP14), neutrophil granule proteins (PRTN3, ELANE, AZU1), and Fc-receptor/NADPH-oxidase components (ITGB2, FCGR1, NCF1, NCF2, NCF4, CYBB), and are therefore best interpreted as a single ECM-remodeling and innate immune program that is more pronounced in younger LS patients.

In MSS, enrichment was limited to two terms among positively age-correlated proteins, anatomical structure morphogenesis (3 proteins) and regulation of cell adhesion (2 proteins), with weak significance (**Figure 5D**); no significant terms were identified among negatively age-correlated MSS proteins. Functional annotation of the MLH1_hm_ age-correlated proteins likewise returned no significant terms in either direction. The limited enrichment in these two subtypes likely reflects the small number of proteins meeting the |r| ≥ 0.5 threshold (42 in MLH1_hm_ and 35 in MSS).

### Sex-Associated Protein Dysregulation in CRC

To determine whether the degree of tumor-specific proteomic dysregulation differs by patient sex, we compared patient-specific tumor-normal protein dysregulation (tumor minus matched normal mucosa, log_2_ scale) between male and female patients within each of the four CRC molecular subtypes using an independent two-sample Student’s t-test, restricted to each subtype’s set of proteins significantly dysregulated between tumor and matched normal tissue (LS: 648 proteins; DS: 1,036; MLH1_hm_: 906; MSS: 604). Within each subtype-specific set, proteins with p < 0.05 were considered sex-associated, and the 10 proteins with the smallest p-values per subtype were selected for visualization (**Figure 6**). This identified 21 sex-associated proteins in LS, 24 in DS, 15 in MLH1_hm_, and 15 in MSS.

Given the sex distribution of each subtype’s cohort (Table 1), the male and female subgroups compared were small and, in most subtypes, highly unequal: approximately 3 female vs. 9 male patients in LS, 5 vs. 4 in DS, 14 vs. 6 in MLH1_hm_, and 8 vs. 12 in MSS. A two-sample t-test comparing groups as small and imbalanced as 3 vs. 9 (LS) is particularly underpowered and sensitive to individual outlier values. No correction for multiple hypothesis testing was applied, and p-value ranking reflects the strength of statistical evidence for a sex difference rather than its magnitude; these considerations are addressed further in the Discussion.

The top 10 sex-associated proteins were largely distinct across CRC subtypes, with KAT7 shared between the DS and MSS top ten sets (Figure 6). In LS, the most significantly sex-associated proteins were MZT2B (p = 0.0048) and RSL24D1 (originally RPL24) (p = 0.0080), followed by RIOX2, ARHGDIB (originally GDIR2), PTPRC, IFI16, RAB11FIP1 (originally RUFY1), WDR36, IPO4, and MTHFD2 (p = 0.0204–0.0345). In DS, YIPF5 (p = 0.0047) and HSPA13 (originally HSPB7) (p = 0.0094) were most significant, followed by CDV3, DEFA1 (originally DEF1), ITGA2, KAT7, H1-5, C1orf174 (originally CA174), EIF4B, and SFSWAP (p = 0.0167–0.0342). In MLH1hm, LIMD2 (p = 0.0099) and LDHA (p = 0.0146) were most significant, followed by CNN2, HELZ, GP1BB, LAP3 (originally AMPL), SUMO1, TMEM263, FUT8, and TRIP13 (p = 0.0152–0.0312). In MSS, KAT7 (p = 0.0045) was most significant, followed by NKRF (p = 0.0081), SERPINB8, FBP1, MGAT2, TBC1D2B, NQO1, NUDT1 (originally OGG1), MESD, and CCDC137 (p = 0.0149–0.0410 for the latter eight proteins). Notably, LDHA also appeared among the age-correlated proteins in the MLH1hm subtype (Figure 4).

This raises the possibility of an interaction between sex and age effects on LDHA dysregulation specifically within MLH1_hm_, though confirming this would require formal statistical testing (e.g., an interaction term or sex-stratified age correlation) rather than being inferred from its appearance in both independent analyses.

Utilizing a subset of the significant sex-associated proteins, samples from the CPTAC-2 cohort were analyzed to compare tumor and normal mucosa samples in response to gender-assigned at birth. Across 18 analyzed proteins, 12 of the proteins saw significance between tumor and normal mucosa in their specific sex-subsets (Figure S10). As not all proteins demonstrated statistical significance, this data provides evidence towards the consideration that there may be subset specific sex-associated markers within the proteomics datasets.

## DISCUSSION

### Global Subtype Differences and Shared Proteomic Differences Across CRC

Across the four subtypes, our proteomic analysis — applying both a Benjamini-Hochberg adjusted p-value < 0.05 and an absolute log_2_ fold-change > 1 threshold — identified 1,607 proteins significantly dysregulated in tumor tissue relative to normal controls in at least one subtype, of which 260 were commonly dysregulated across all four subtypes (Figure 2A), suggesting a core set of proteomic alterations underpinning colorectal tumorigenesis regardless of mismatch repair or microsatellite status. Functionally, this shared dysregulated proteome reflects fundamental hallmarks of CRC biology. Consistent with this, GO enrichment of each subtype’s dysregulated proteins identified ribosome biogenesis and RNA processing as the top-enriched processes in every subtype, and DNA replication, extracellular matrix organization, and muscle system processes were enriched in all four (**Figure 3A, Supplemental Figures 4– 6**), consistent with the proliferative and biosynthetic demands of cancer cells² and with remodeling of the tumor stroma.^34^ This is in line with prior proteomic studies showing elevated abundance of gene expression machinery and replication factors in CRC.^34^

Hierarchical clustering of the 260 shared proteins by log₂ fold-change (**Figure 2B**) separated MSS from the three MMR-deficient subtypes, which clustered together despite arising through germline (LS), epigenetic (MLH1_hm_), and biallelic somatic (DS) routes. This suggests that MMR status itself, rather than the mechanism of MMR loss, is the dominant driver of the shared proteomic signature.

Comparing fold-changes directly across subtypes (**Table 2**) further showed that most of the proteins strongly altered in LS are dysregulated in the same direction in all four subtypes, differing only in magnitude. HKDC1 was the clear exception: upregulated in LS but not significantly changed in any other subtype. HKDC1 also emerged independently in the GO analysis of LS-unique proteins (**Figure 3B**), where it contributed to a glycolytic signature together with LDHA and PFKFB3 and the bicarbonate transporter SLC4A4. Glycolytic terms were not enriched in any other subtype, either across all dysregulated proteins or among subtype-unique proteins (**Supplemental Figures S4–S6**). HKDC1 is highly expressed in CRC relative to other cancers and promotes CRC cell proliferation and glycolysis, making it a candidate LS-distinctive metabolic alteration that merits further investigation. ^35-37^

### Ribosome Biogenesis and DNA Replication, Not DNA Repair

Ribosome biogenesis dominated the LS enrichment results, both among all dysregulated LS proteins (**Figure 3A**) and among those unique to LS (**Figure 3B**). Elevated ribosome production is a recognized hallmark of cancer cells, yet it has received comparatively little attention in CRC, and much of the existing evidence comes from cell lines, organoids, and mouse models. ^38^ Our data provides complementary evidence from primary human tumors.

Given the centrality of MMR deficiency to LS, we specifically examined the enrichment results for DNA repair. No term related to mismatch repair, DNA damage response, homologous recombination, or excision repair was enriched; the DNA-related signal reflected replication machinery (including MCM2/4/5/6/7, PCNA, RFC4/5, TOP1, RRM1, and CHAF1A/B) rather than repair. The same was true across subtypes: no mismatch repair term was enriched in any of the four, and the only repair-related term, base-excision repair in DS, was of borderline significance and shared several components with replication machinery (PCNA, LIG1, RPA3) (Figure S4). The absence of an MMR signal should not be read as intact MMR expression. MMR genes follow Knudson’s two-hit model, in which non-neoplastic mucosa in LS carriers retains expression from the wild-type allele, and the corresponding protein is typically lost in the tumor after a second somatic hit, ^39^ therefore a tumor-versus-normal decrease would be expected. Two features of our data likely obscured it. MLH1 and PMS2, the proteins most frequently lost in LS tumors, were not quantified, and our LS cohort includes carriers of different MMR gene variants, thus loss of any single protein would be confined to a subset of patients and diluted in a group comparison. Consistent with this, MSH2 and MSH3 trended downward in LS without reaching significance. MSH6 was significantly upregulated in MLH1_hm_ and MSS, possibly reflecting a compensatory response to mismatch accumulation, though this requires further investigation.

### Age-Associated Proteomic Signatures in CRC

The proportion of dysregulated proteins correlated with age was highest in LS and DS, the two smallest cohorts. As smaller sample subsets make it easier for a correlation to exceed a fixed effect-size threshold by chance, we regard these findings as hypothesis-generating. The strongest individual correlations (Figure 4), such as the decline in FCN1 and BUD23 dysregulation with age in LS, were nonetheless clearly visible at the level of individual patients (Figure S7).

In LS, proteins whose dysregulation was greater in younger patients were enriched for extracellular matrix remodeling, hypoxia response, and innate immune signaling. The "osteoclast differentiation" term largely reflects Fc-receptor and NADPH-oxidase innate immune components in this context within KEGG rather than bone biology. This is broadly consistent with reports that early-onset CRC is clinically more aggressive^40^ and carries a distinct, proinflammatory innate immune signature, ^41^ although the immune composition of early-onset tumors remains debated, ^42^ and these studies compare age groups across CRC broadly rather than within LS.

In DS, the proliferative signature (ribosome biogenesis, rRNA processing, spliceosome) intensified with age, which, to our knowledge, has not previously been reported and is presented as a novel observation. Conversely, signal transduction, Notch, interleukin-1, and stem cell proliferation pathways were more prominent in younger DS patients, paralleling the elevated growth-factor and TNFR1 signaling reported in sporadic early-onset relative to older-onset MSS CRC, ^40^ a related but molecularly distinct subtype. Age-associated functional enrichment was minimal in MSS and absent in MLH1_hm_, and PPHLN1’s opposing age relationships across subtypes indicate that some age effects are pathway-specific rather than generic features of CRC.

Two features of the cohort constrain interpretation. First, the subtypes occupy largely non-overlapping age ranges (**Figure S7**), consistent with the earlier onset of LS and the later onset of MLH1_hm_ CRC, ^33^ where each subtype’s correlations therefore describe a different portion of the lifespan and should not be compared directly across subtypes. Second, because only age at diagnosis was available, we cannot separate biological aging from differences in time between tumor onset and clinical presentation.

### Subtype-Specific Sex-Associated Differences

Sex-associated proteins were largely subtype-specific (**Figure 6**), but functional enrichment of these sets yielded only weakly supported terms and no interpretable pathway signature. Given the small and unbalanced male and female subgroups, particularly in LS, we do not draw functional conclusions from this analysis.

### Limitations

Several methodological choices warrant consideration. First, age-associated proteins were selected by an effect-size threshold (|r| ≥ 0.5) rather than a p-value criterion because cohorts of 9–20 patients are underpowered for correlation significance testing; the higher proportion of selected proteins in the smaller LS and DS cohorts means these findings require validation in larger cohorts. Second, DAVID annotation of the age-correlated proteins used nominal, uncorrected p-values and should be considered exploratory. Third, the sex-associated analysis compared small, unequal subgroups. Finally, because age at diagnosis rather than age at tumor onset was recorded, apparent age effects may partly reflect time to presentation.

## Conclusions

This study presents, to our knowledge, the first large-scale proteomic comparison of Lynch Syndrome colorectal cancer with Double Somatic, MLH1-hypermethylated, and microsatellite-stable tumors, using paired tumor and matched normal mucosa from 61 patients. We identified 1,607 proteins dysregulated in at least one subtype, including a core set of 260 shared across all four, reflecting conserved features of colorectal tumorigenesis such as enhanced ribosome biogenesis, RNA processing, and DNA replication alongside loss of epithelial differentiation markers. Within this shared signature, the three mismatch repair-deficient subtypes clustered apart from MSS tumors, suggesting that MMR status, rather than the mechanism by which it is lost, shapes the shared proteomic landscape. Beyond these shared changes, LS tumors showed a distinct profile characterized by ribosome biogenesis and a glycolytic signature, with HKDC1 emerging as a candidate LS-distinctive alteration. Exploratory analyses further suggested that tumor-associated dysregulation varies with age at diagnosis in a subtype-specific manner, with extracellular matrix remodeling and innate immune signaling more pronounced in younger LS patients. Given the modest cohort sizes, particularly for LS and DS, these age- and sex-associated findings should be regarded as hypothesis-generating, and validation in larger, independent cohorts will be essential, including stratification by the specific MMR gene affected in LS. Together, these data provide a resource for understanding the proteomic basis of hereditary versus sporadic colorectal cancer and for identifying candidate protein markers relevant to LS, and the complete dataset has been made available through the PRIDE repository (PXD073693).

## Data Availability

The complete dataset has been made available through the PRIDE repository (PXD073693).

## Current Addresses

Fernando Tobias; Agilent Technologies, Inc., 201 Hansen Ct #108 Wood Dale, IL 60191, USA

Emily R. Sekera: Department of Immunology, St. Jude Children’s Research Hospital, Memphis, TN, USA, 262 Danny Thomas Pl., Memphis, TN 38105

Xingzhao Xiong: Division of Biomedical Informatics and Genomics, John W. Deming Department of Medicine, 1441 Canal St, New Orleans, LA 70112, USA

Rachel Pearlman: The Ohio State University Comprehensive Cancer Center, 2012 Kenny Road, Columbus, OH 43221, USA

Heather Hampel: Department of Medical Oncology and Therapeutics Research and Division of Clinical Cancer Genomics, City of Hope, Duarte Cancer Center, Duarte, CA 91010, USA

Xiaowen Liu: Division of Biomedical Informatics and Genomics, John W. Deming Department of Medicine, 1441 Canal St, New Orleans, LA 70112, USA

Liangliang Sun: Department of Chemistry, Michigan State University, East Lansing, MI 48824, USA

Amanda B. Hummon: Department of Chemistry and Biochemistry and the Comprehensive Cancer Center, The Ohio State University, Columbus, Ohio, 43210, USA

## Author Contributions

FT, ERS, XX, FF, HH, RP, XL, LS, and ABH designed the study

FT and ERS collected the mass spectrometry data

FT, ERS, XX, FF, and XL analyzed the data

FT, ERS, XX, FF, HH, RP, XL, LS, and ABH wrote the manuscript

## Acknowledgements

The Ohio Colorectal Cancer Prevention Initiative (OCCPI) was supported by a grant from Pelotonia, an annual cycling event in Columbus, Ohio that supports cancer research at The Ohio State University Comprehensive Cancer Center – James Cancer Hospital and Solove Research Institute. The OCCPI was also supported in part by a grant P30 CA016058, National Cancer Institute, Bethesda, MD. Additionally, the tims-ToF Pro utilized within this study was purchased using funds from NIH award S10 OD026945.

RP, XL, LS and ABH were funded by NIH through the grant R01CA247863. ABH is also grateful for support of R35GM158423. FF and LS thank the support from the National Institute of General Medical Sciences (NIGMS) through Grant R35GM153479. We also thank MSU AgBioResearch and the Michigan State University for access to the QIAGEN IPA platform. FT and ABH were funded by R21AG062144. ERS and ABH were funded through R01GM110406 and ABH is funded through R35GM158423.

Additional thanks go to Dr. Suresh Poudel of St. Jude Children’s Research Hospital for assistance in revisions of the updated manuscript.

## Supporting Information

**Table S1:** Significant proteins detected in each cohort. The data is available as an Excel file with separate tabs for each sample subtype and a tab with the 260 dysregulated proteins.

**Table S2:** Gene Ontology Biological Process enrichment results.

**Table S3:** Parameters used in the DIA-NN search. The data is available as an Excel file.

**Table S4**: Complete MSStats results. The data is available as an Excel file with separate tabs for the protein-level abundances and the full MSStats differentials results for each subtype.

**Figure S1:** Distribution of log_2_ fold-changes across quantified proteins in each CRC subtype.

**Figure S2:** Comparison of top 10 downregulated proteins in Lynch Syndrome to TCGA-COAD gene expression results.

**Figure S3:** Comparison of top 10 upregulated proteins in Lynch Syndrome to TCGA-COAD gene expression results.

**Figure S4:** Top Biological Process (BP) terms from Gene Ontology (GO) analysis for the Microsatellite Stable (MSS) subtype.

**Figure S5:** Top Biological Process (BP) terms from Gene Ontology (GO) analysis for the MLH1 Hypermethylation (MLH1_hm_) subtype.

**Figure S6:** Top Biological Process (BP) terms from Gene Ontology (GO) analysis for the Double Somatic (DS) subtype.

**Figure S7:** Subtype-specific age-associated protein expression trends in colorectal cancer.

**Figure S8:** Comparison of top 10 downregulated proteins in Lynch Syndrome to CPTAC-2 protein expressions.

**Figure S9**: Comparison of top 10 upregulated proteins in Lynch Syndrome to CPTAC-2 protein expressions.

**Figure S10:** Protein expression analysis comparison from CPTAC-2 data for select proteins from sex-associated analyses.

