## Supplemental Figures for "Global Proteomic Analysis of Colorectal Cancers Stratified by Microsatellite Instability Subtype Reveals Protein Differences"

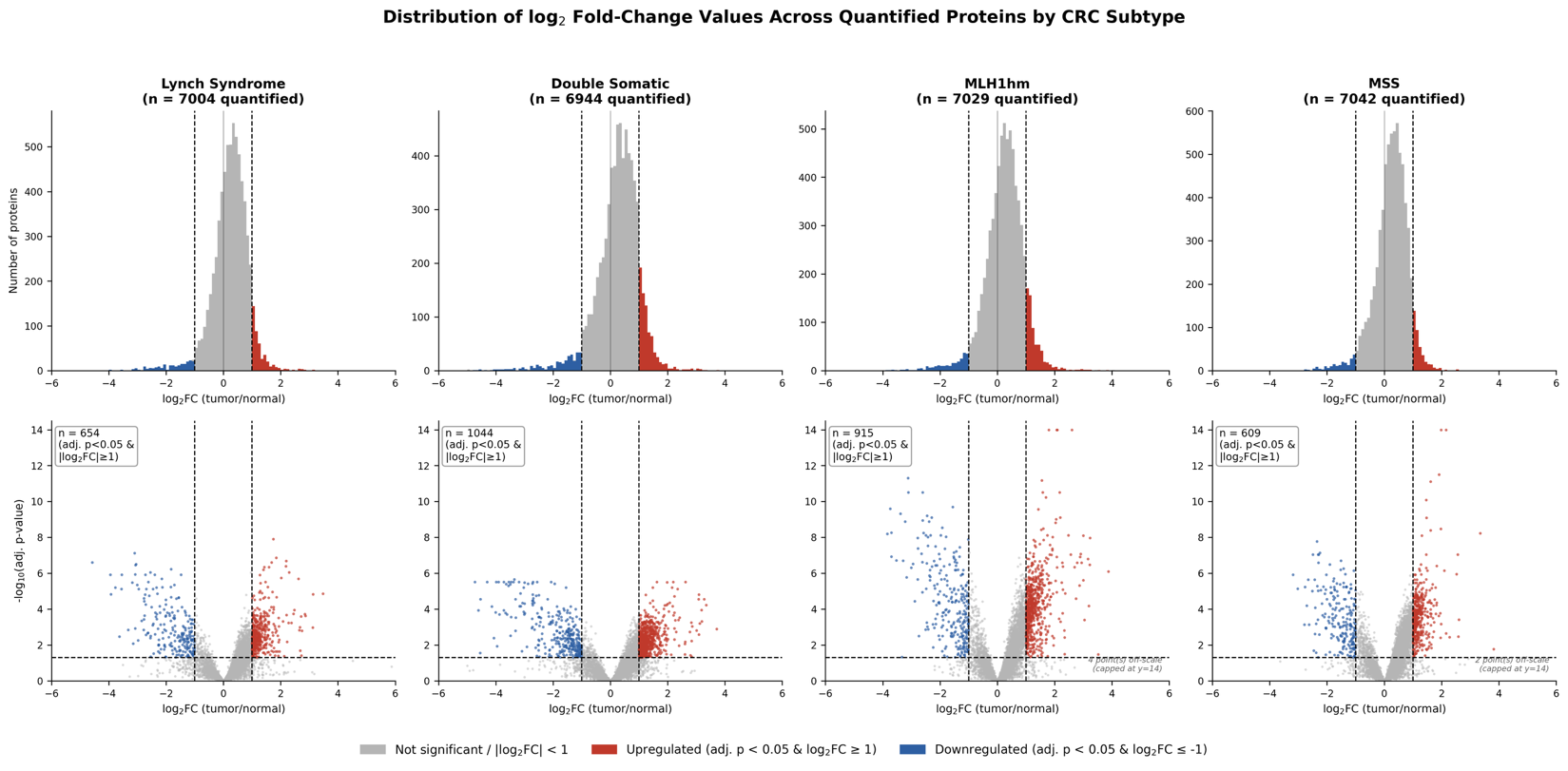

**Figure S1: Distribution of log₂ fold-changes across quantified proteins in each CRC subtype.** Top row: histograms of log₂ fold-change (tumor vs. matched normal mucosa) for all proteins with a finite fold-change and an adjusted p-value in Lynch Syndrome (LS), Double Somatic (DS), MLH1 Hypermethylation (MLH1hm), and Microsatellite Stable (MSS) tumors. Bottom row: corresponding volcano plots of log₂ fold-change against −log₁₀(Benjamini-Hochberg adjusted p-value). Proteins meeting both significance criteria (adjusted p < 0.05 and |log₂FC| ≥ 1) are shown in red (upregulated) or blue (downregulated); all others are shown in gray. Dashed lines mark the thresholds (log₂FC = ±1; adjusted p = 0.05). A small number of extreme values are capped at −log₁₀(adjusted p) = 14 for display. Proteins detected in only one tissue type, which have an infinite fold-change, are excluded. Counts of significant proteins shown here (654, 1,044, 915, and 609) differ by fewer than ten per subtype from the main-text totals (648, 1,036, 906, and 604) because a small number of proteins were excluded from downstream analyses due to ambiguous or missing gene-symbol mapping.

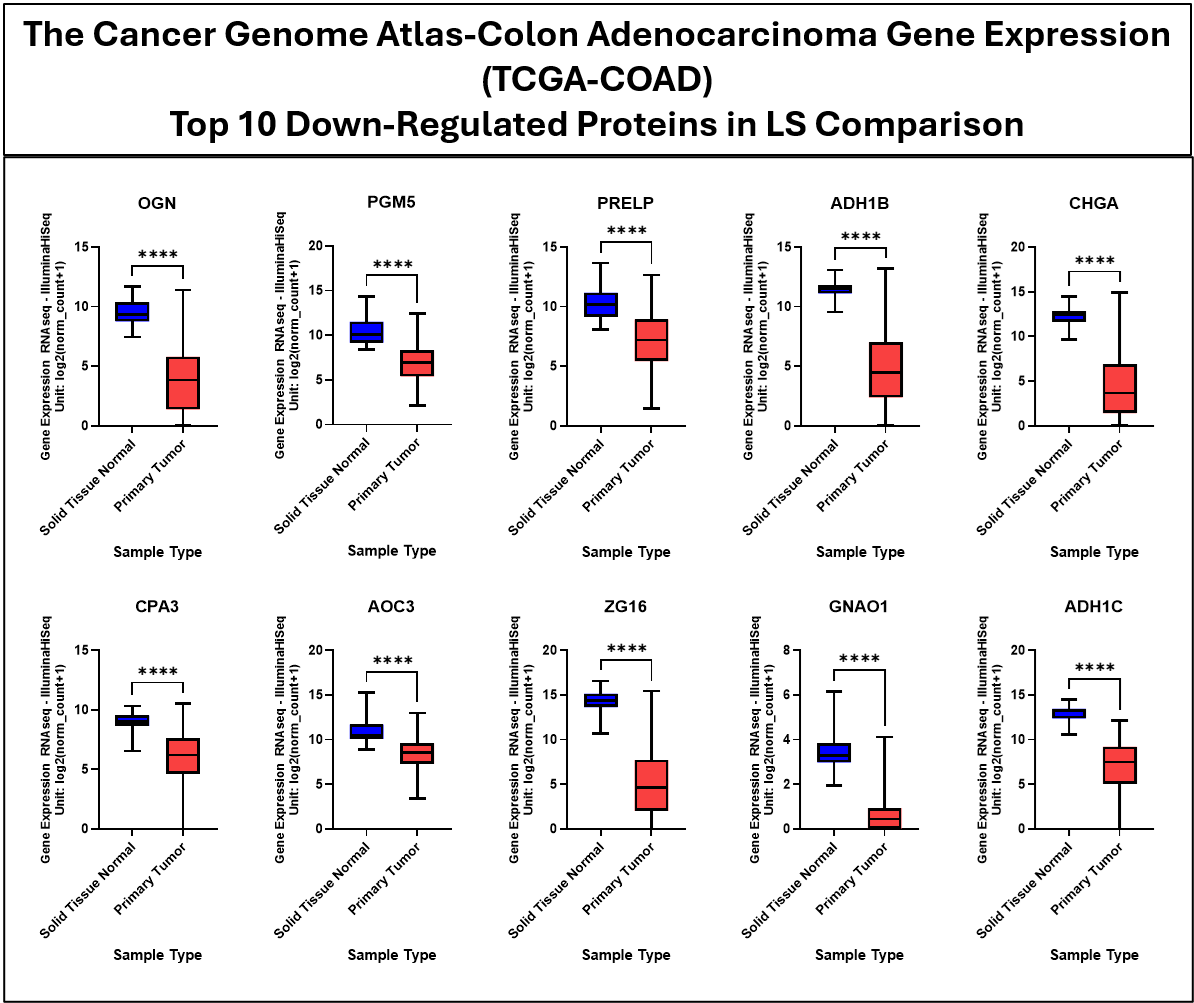

**Figure S2: Gene Expression Analysis Data from TCGA-COAD for Top 10 Downregulated Proteins Observed in LS Dataset.** The provided box and whisker plots show the gene expression data from the TCGA-COAD cohort comparing solid normal tissue (n= 41) to primary tumor tissues (n=286). Significance values are denoted as follows: ns p > 0.05, * p ≤ 0.05, ** p ≤ 0.01, *** p ≤ 0.001, **** p ≤ 0.0001.

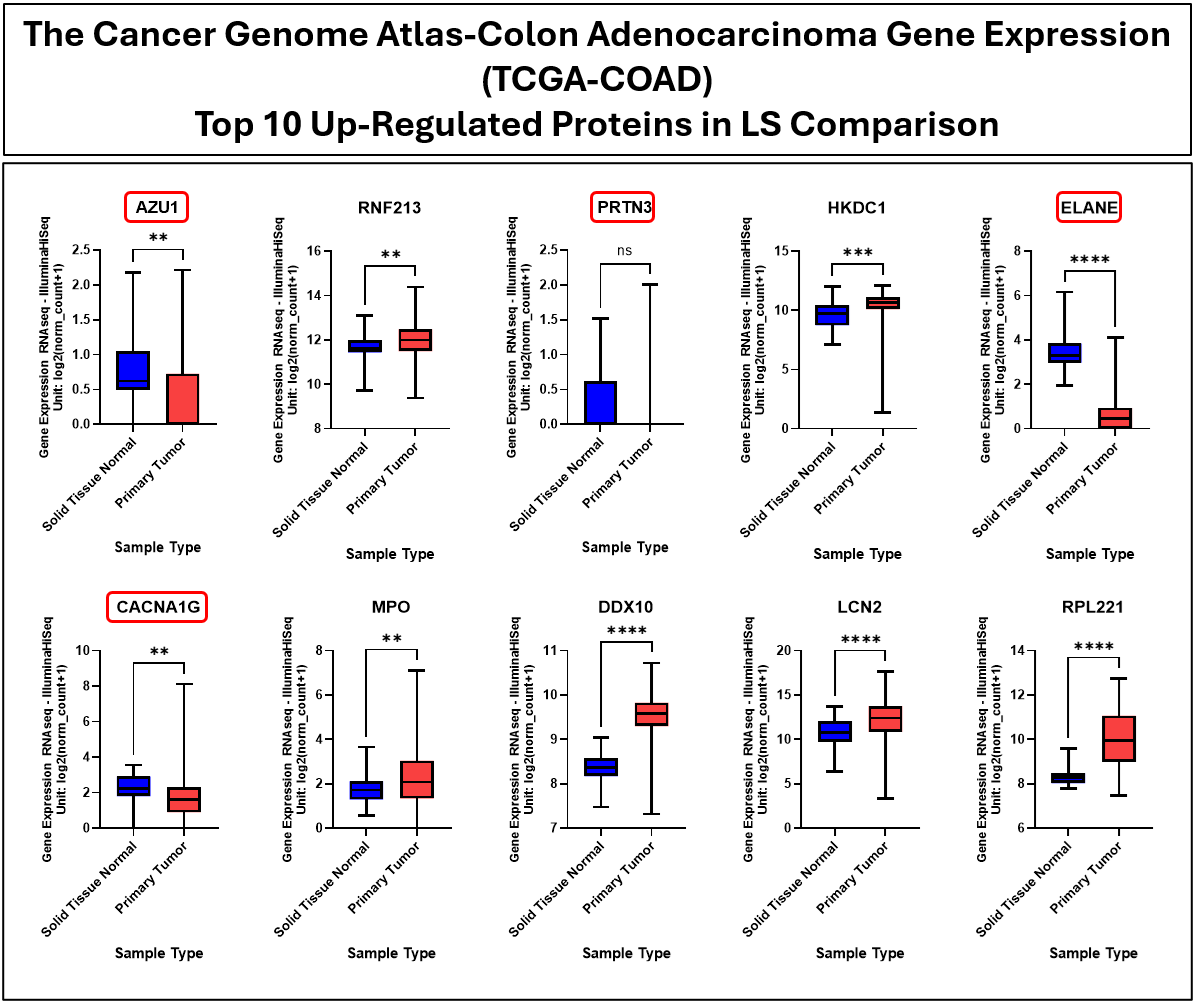

**Figure S3: Gene Expression Analysis Data from TCGA-COAD for Top 10 Upregulated Proteins Observed in LS Dataset.** The provided box and whisker plots show the gene expression data from the TCGA-COAD cohort comparing solid normal tissue (n= 41) to primary tumor tissues (n=286). Gene names boxed in red denote genes that do not match with the data observed in the discussed proteomics analyses. Significance values are denoted as follows: ns p > 0.05, * p ≤ 0.05, ** p ≤ 0.01, *** p ≤ 0.001, **** p ≤ 0.0001.

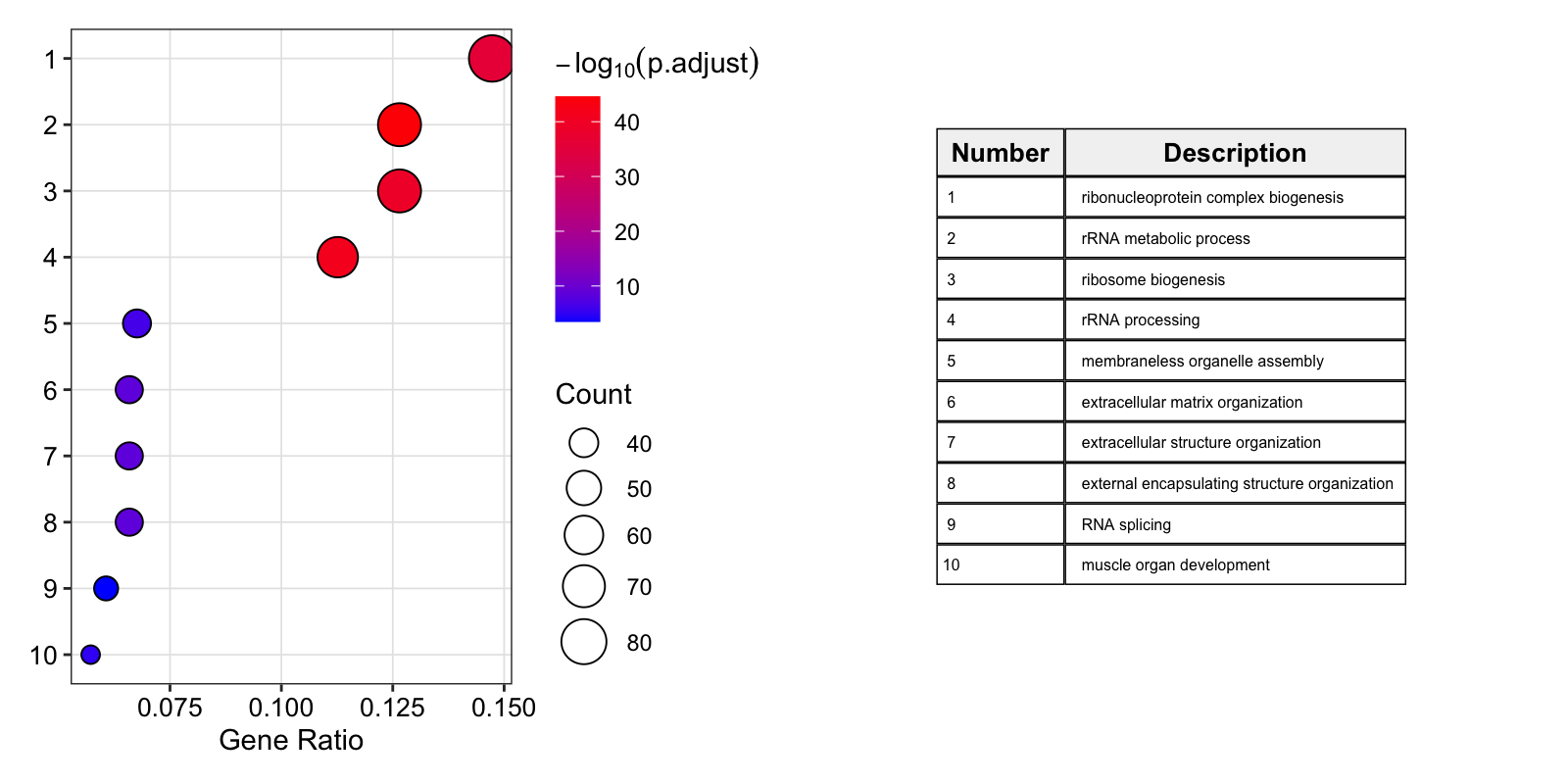

**Figure S4:** **Gene Ontology Biological Process enrichment for the Microsatellite Stable (MSS) subtype.** Top 10 enriched terms among the 604 proteins dysregulated in MSS tumors relative to matched normal mucosa. Over-representation analysis with clusterProfiler (Benjamini-Hochberg adjusted p-value < 0.05, q-value < 0.2, human genome background). Terms are ranked by gene ratio; bubble size indicates the number of proteins annotated to each term, and color indicates −log_10_(adjusted p-value). Analysis of the 134 MSS-unique proteins yielded no significant terms and is not shown.

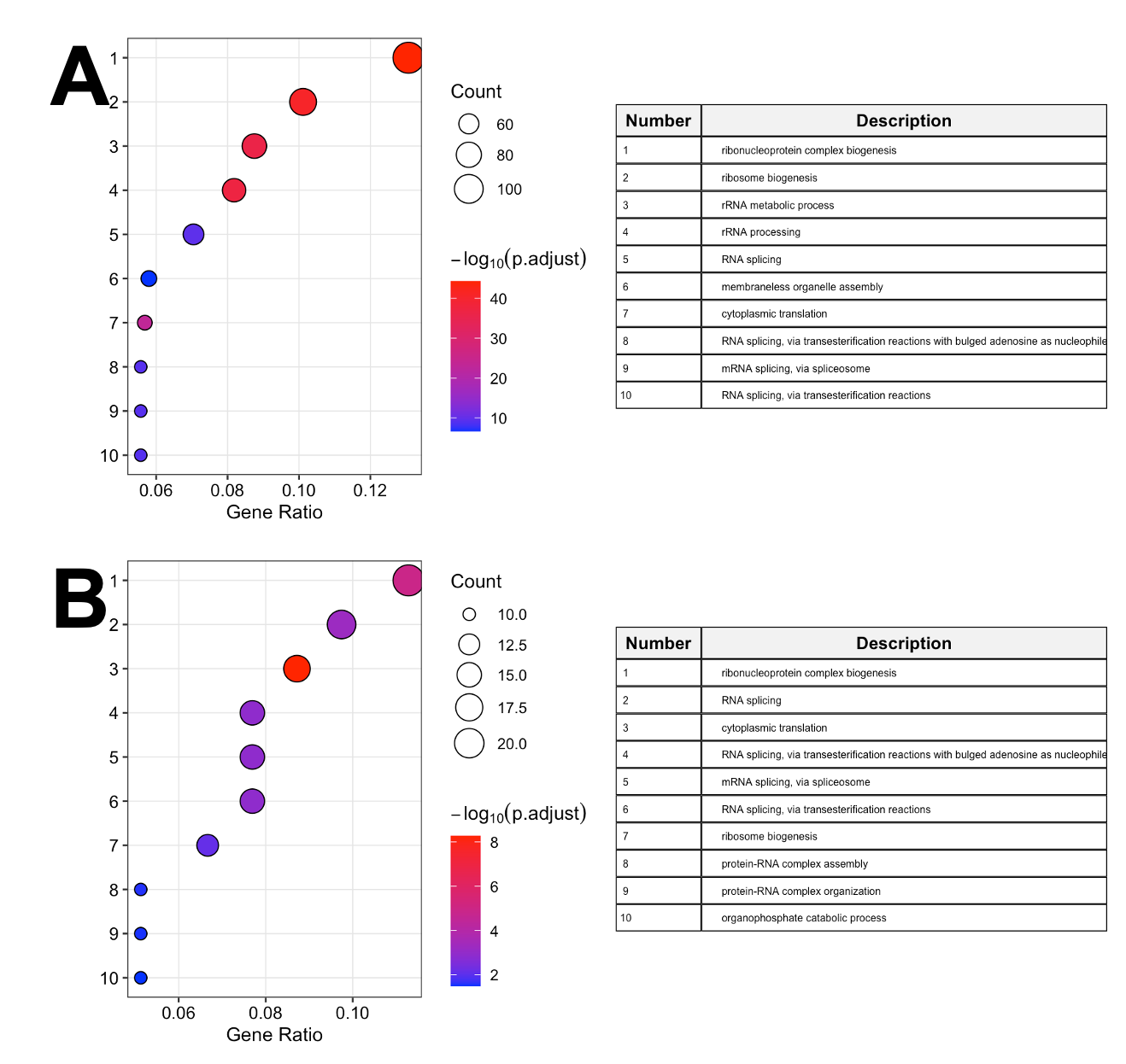

**Figure S5: Gene Ontology Biological Process enrichment for the MLH1 Hypermethylation (MLH1_hm_) subtype.** Analysis as in Figure S1. (A) Top 10 enriched terms among the 906 proteins dysregulated in MLH1_hm_ tumors relative to matched normal mucosa. (B) Top 10 enriched terms among the 203 proteins dysregulated uniquely in MLH1_hm_.

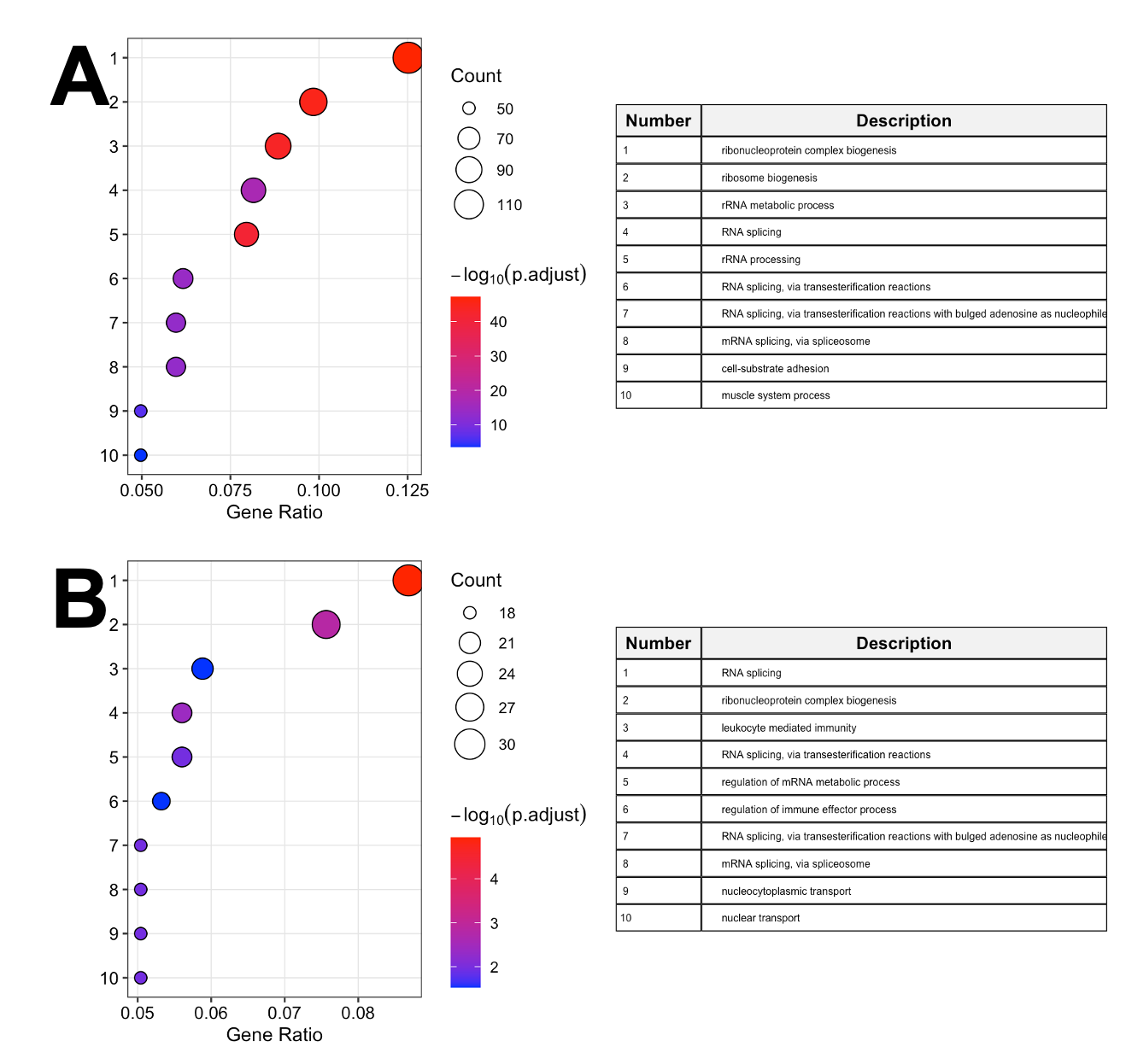

**Figure S6:** **Gene Ontology Biological Process enrichment for the Double Somatic (DS) subtype.** Analysis as in Figure S1. (A) Top 10 enriched terms among the 1,036 proteins dysregulated in DS tumors relative to matched normal mucosa. (B) Top 10 enriched terms among the 374 proteins dysregulated uniquely in DS.

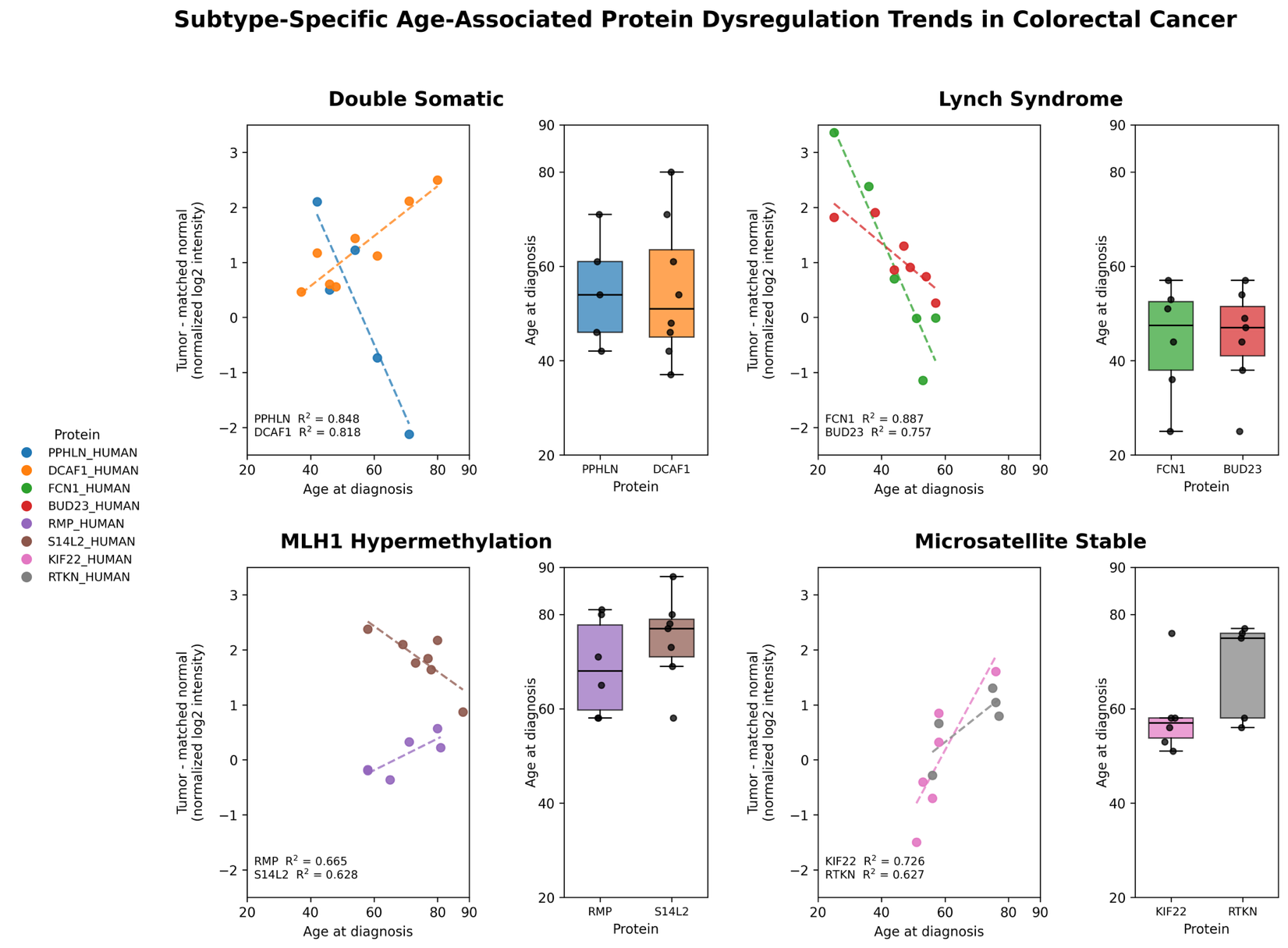

**Figure S7: Relationship between age at diagnosis and tumor-normal protein dysregulation for the top age-associated proteins in each CRC subtype.** For each subtype, the two proteins with the largest absolute Pearson correlation (|r|) from Figure 4 are shown: Double Somatic (DS), PPHLN1 and DCAF1; Lynch Syndrome (LS), FCN1 and BUD23; MLH1 Hypermethylation (MLH1_hm_), RPRD1B (RMP) and SEC14L2 (S14L2); and Microsatellite Stable (MSS), KIF22 and RTKN. Scatter plots show patient-specific protein dysregulation, calculated as normalized log₂ intensity in tumor minus that in the matched normal mucosa from the same patient, against age at diagnosis. Each point represents one patient, dashed lines indicate linear regression fits, and R² values are shown for each protein. Adjacent boxplots show the age at diagnosis of the patients contributing to each protein's trend (boxes indicate the median and interquartile range; points indicate individual patients). The number of points per protein can be lower than the subtype cohort size because not every protein was quantified in both tumor and normal tissue for every patient, and because patients diagnosed at the same age (present in LS, MLH1hm, and MSS) may overlap.

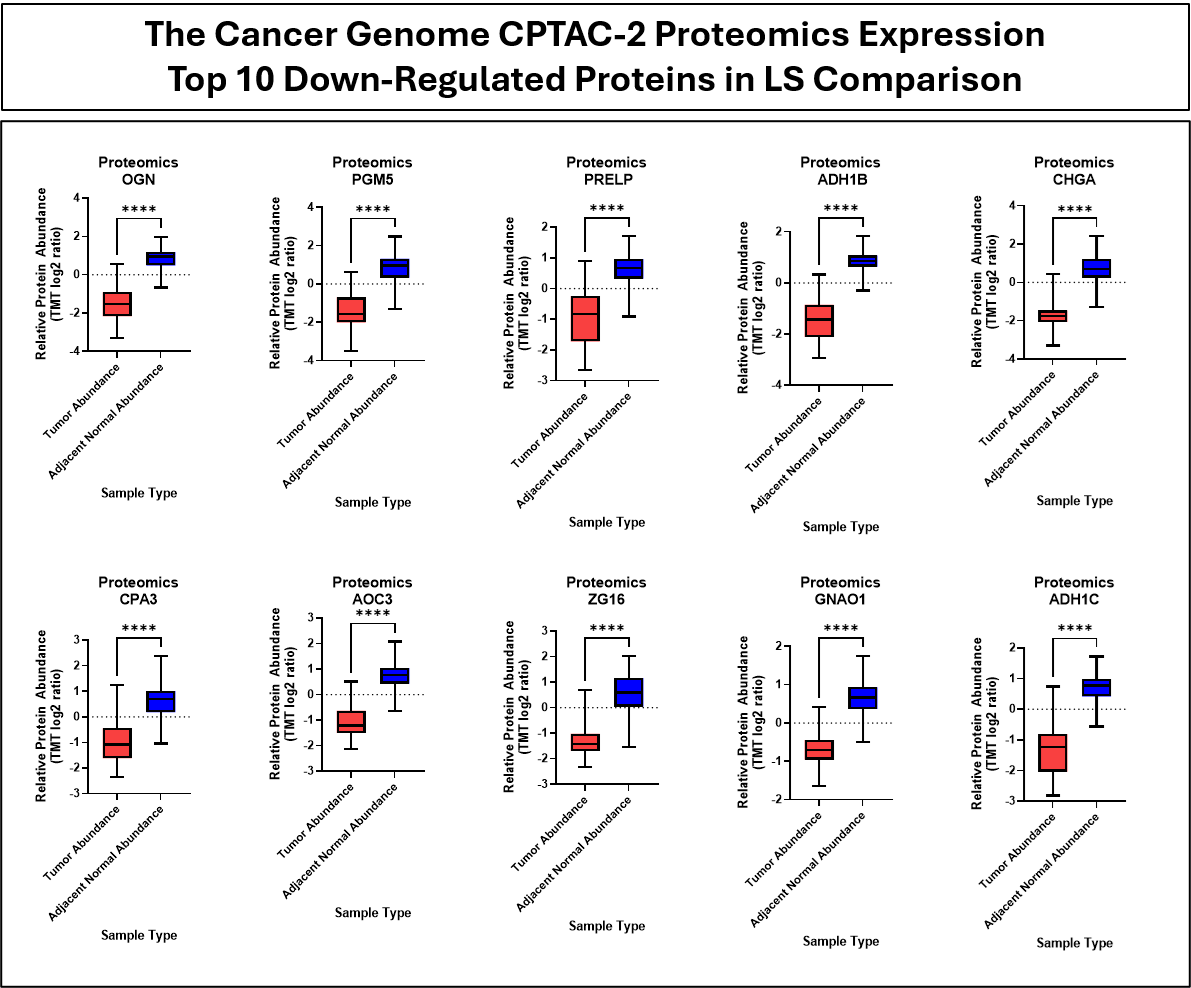

**Figure S8: Protein Analysis Data from CPTAC-2 for Top 10 Downregulated Proteins Observed in LS Dataset.** The provided box and whisker plots show the protein expression data from the CPTAC-2 colorectal cancer cohort comparing solid normal tissue to primary tumor tissues (patient sample size n=101, protein expression is not observed in every sample). Significance values are denoted as follows: ns p > 0.05, * p ≤ 0.05, ** p ≤ 0.01, *** p ≤ 0.001, **** p ≤ 0.0001.

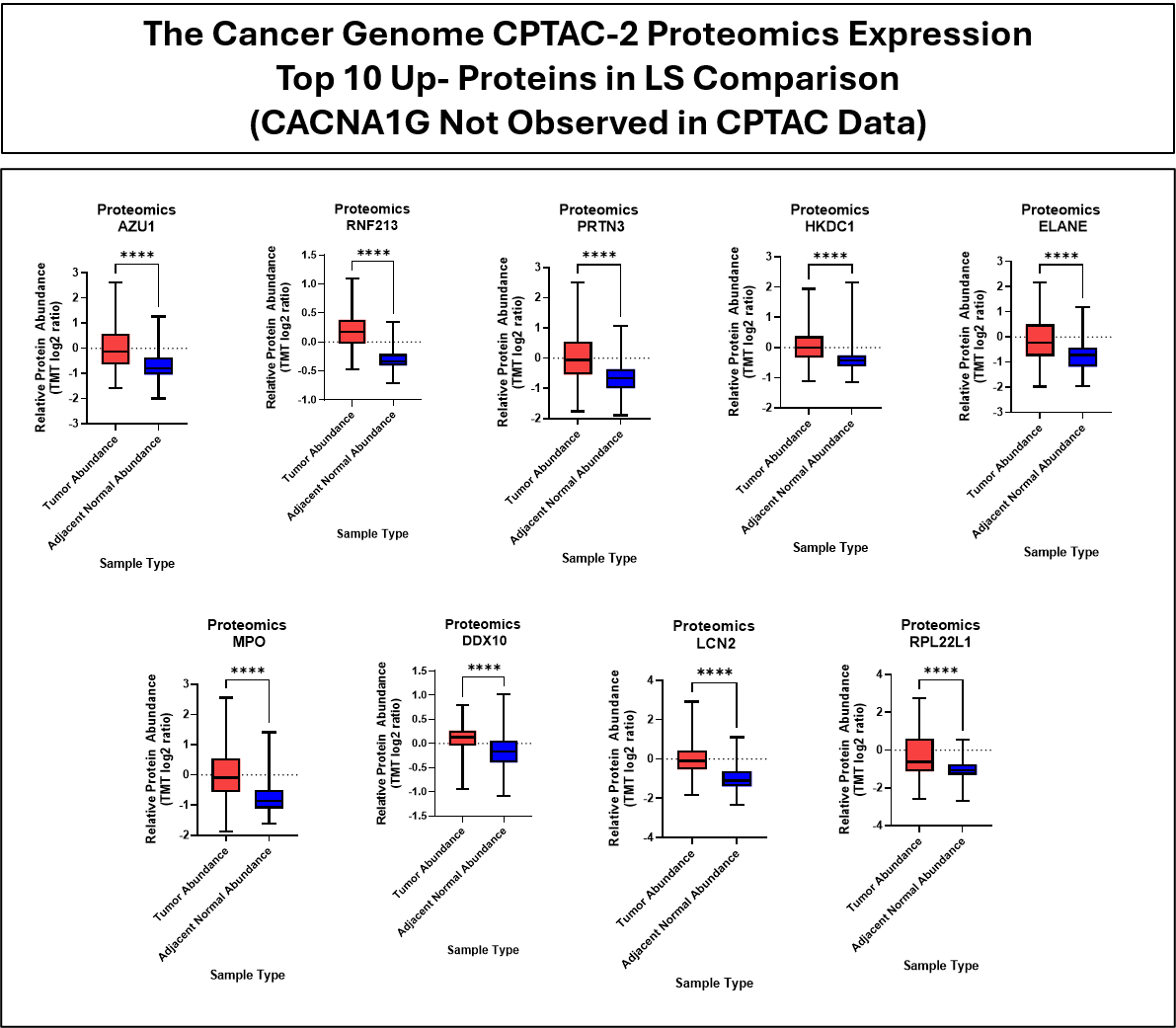

**Figure S9: Protein Analysis Data from CPTAC-2 for Top 9 Upregulated Proteins Observed in LS Dataset.** The provided box and whisker plots show the protein expression data from the CPTAC-2 colorectal cancer cohort comparing solid normal tissue to primary tumor tissues (patient sample size n=101, protein expression is not observed in every sample). Significance values are denoted as follows: ns p > 0.05, * p ≤ 0.05, ** p ≤ 0.01, *** p ≤ 0.001, **** p ≤ 0.0001.

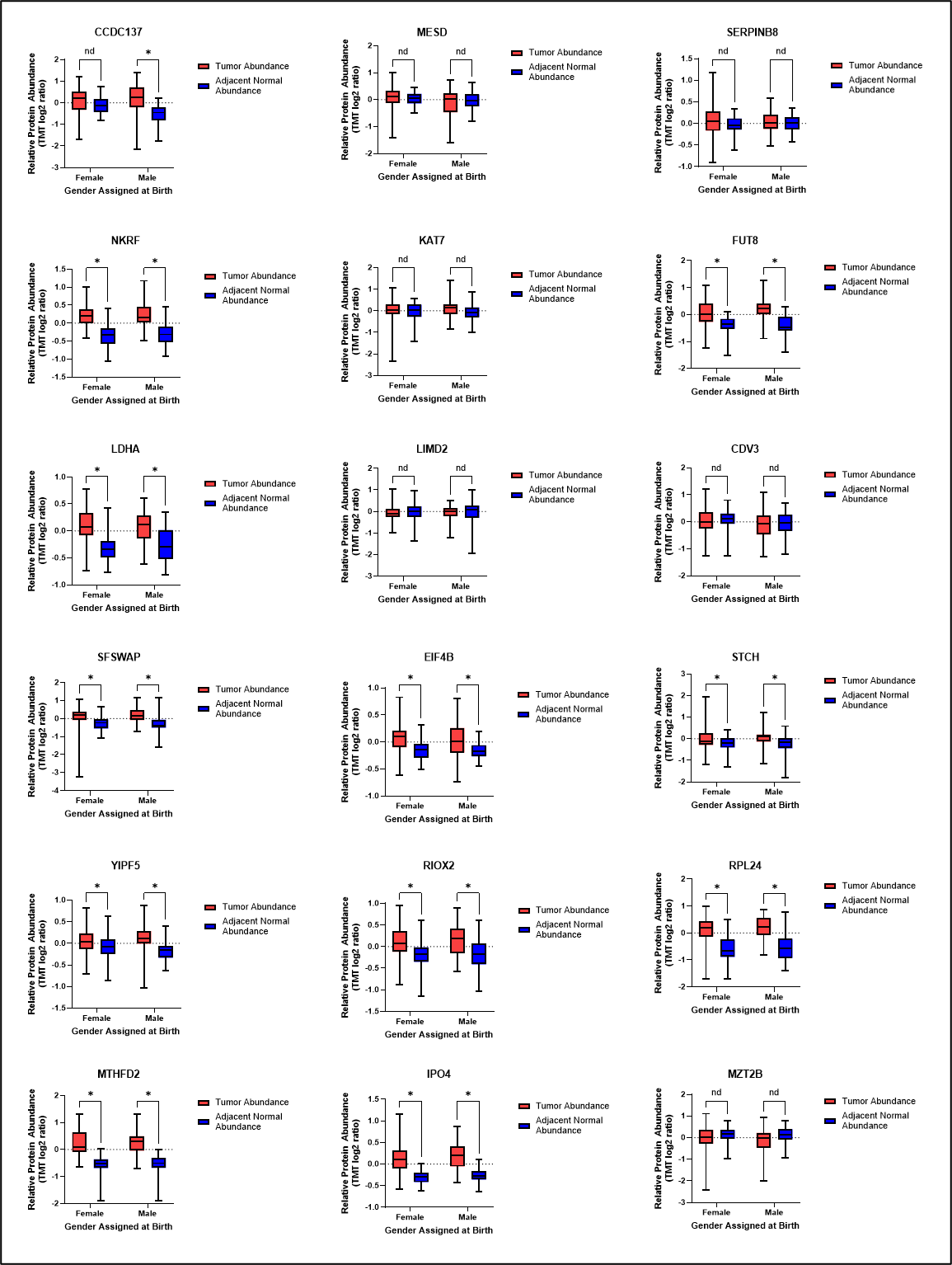

**Figure S10: Protein Expression Analysis for Select Proteins from Sex-Associated Analyses.** The top and bottom two proteins listed in Figure 6 were chosen as well as any overlapping proteins to evaluate statistical significance occurring between tumor and adjacent normal tissues separated by sex. Significance values are denoted as follows: * p ≤ 0.05.
